# Dissociable mechanisms govern when and how strongly reward attributes affect decisions

**DOI:** 10.1101/434860

**Authors:** Silvia U. Maier, Anjali Raja Beharelle, Rafael Polanía, Christian C. Ruff, Todd A. Hare

**Affiliations:** Zurich Center for Neuroeconomics, Department of Economics, University of Zurich, Zurich, Switzerland; Neuroscience Center Zurich, University of Zurich, Swiss Federal Institute of Technology Zurich, Zurich, Switzerland; Translational Neuromodeling Unit, Institute for Biomedical Engineering, University of Zurich and ETH Zurich, Zurich, Switzerland; Decision Neuroscience Lab, Department of Health Sciences and Technology, ETH, Swiss Federal Institute of Technology, Zurich, Switzerland

## Abstract

Theories and computational models of decision making usually focus on how strongly different attributes are weighted in choice, e.g., as a function of their importance or salience to the decision-maker. However, when different attributes impact on the decision process is a question that has received far less attention. Here, we investigated whether attribute consideration timing has a unique influence on decision making using a time-varying drift diffusion model and data from four separate experiments. Experimental manipulations of attention and neural activity demonstrated that we can dissociate the processes that determine the relative weighting strength and timing of attribute consideration. Thus, the processes determining either the weighting strengths or the timing of attributes in decision making can adapt independently to changes in the environment or goals. Quantifying these separate influences of timing and weighting on choice improves our understanding and predictions of individual differences in decision behaviour.

## Introduction

Decisions regularly involve comparisons of several attributes of the choice options. Consider the example of deciding between foods that differ in two attributes, tastiness and healthiness. Often these attributes are misaligned, creating a conflict between the goal of eating healthfully and the desire to experience pleasant tastes. Typically, we assume that choices for the healthier or better tasting food are determined by the values of these attributes, together with a subjective decision weight that the decision maker assigns to healthiness and taste. The assumption that reward attributes are subjectively weighted in the course of decision making applies not only to food choices, but also to many other types of decisions. In fact, it is a core feature of the standard analysis approaches for intertemporal, social, and risky decisions ^1–4^. Here, we show that this common approach is incomplete because it overlooks the possibility that reward attributes can enter into the decision process at different times (in addition to having different weighting strengths). Across several food choice paradigms, we find that there is considerable asynchrony in when tastiness and healthiness attributes enter into consideration. Furthermore, we demonstrate that the relative weighting strengths (i.e., the degree to which an attribute influences the evidence accumulation rate) and the onset times for tastiness and healthiness attributes in the decision process have separable influences on whether or not people choose to eat healthier foods.

We used an adapted time-varying sequential sampling model that allows for separate attribute consideration onset times to better understand the dynamic decision processes underlying choices between rewards with multiple attributes. This model allows us to draw inferences on latent aspects of the decision process from the observable choice outcomes and response times. It is well established that direct measures and estimates of information acquisition, evaluation, and comparison processes during choice provide a key means of testing predictions from different models of how stimulus and decision values are constructed or used. Uncovering such features of the decision process allows us to discriminate between and evaluate the plausibility of different models that seek to explain choice behaviour ^5^. For example, choice models utilizing not only decision outcomes but also response times and eye- or mouse-tracking data have provided insights into how and why decision-making is influenced by visual attention, time delays or pressure, additional alternatives, and earlier versus later occurring external evidence ^6–13^. Moreover, it has been shown that dynamic accumulation models utilizing response-time data provide a deeper understanding of decisions and make better out-of-sample predictions than reduced form models such as logistic regressions ^14, 15^. Here, we show that we can also use response-time data to determine when specific attributes enter into the decision process, in addition to how strongly they influence the evidence accumulation rate. Moreover, incorporating this information into the model improves predictions about individual decision-making behaviour.

An important implication of the finding that different attributes can enter into the choice process at separate times is that coefficients from traditional regression models (e.g., linear, logit, or probit) will represent a combination of both the true underlying weight or importance placed on each attribute and its relative (dis)advantage in processing time over the decision period. Therefore, any form of static or synchronous onset dynamic model will fail to fully capture the true underlying choice generating process. By static we mean models that treat values or value-differences as fixed rather than being actively constructed. As a consequence, even though such models may explain multi-attribute choice patterns relatively well if the relationship between attribute weighting and timing is fixed or sufficiently stable, they will fail to explain or predict alterations in decision behaviour if attribute weights and processing onset times can change independently in response to external environmental features or changes in internal cognitive strategies. The plausibility of this latter scenario is underlined by mouse-tracking experiments ^16, 17^ showing that different attributes (taste, healthiness) of the same food reward can enter into the decision process at separate times. However, the fundamental question of whether the relationship between attribute weighting strength and timing is stable or instead flexible and context-dependent has not yet been addressed. We addressed this question using an adapted sequential sampling model that quantifies both the weight given to each attribute and its temporal onset during the decision process. This allows us to explicitly measure whether the weighting strength and timing with which different attributes impact on choice are determined by a unitary process (or a set of consistently linked processes), or if, instead, attribute timing and weighting are the results of separable processes. By modelling choices from four separate datasets, which measured decision behaviour under different experimental manipulations (Figure 1), we show that attribute timing and weighting are determined by dissociable decision mechanisms. For example, we find that explicitly instructing individuals to consider either tastiness or healthiness during the choice process ^18^ exerts separate effects on attribute weighting strength and timing. In another experiment, we show that transcranial direct current stimulation (tDCS) over the left dlPFC during food decisions has a selective effect on attribute weighting strength but not timing, demonstrating the separability of the underlying neural processes.

**Figure 1.**
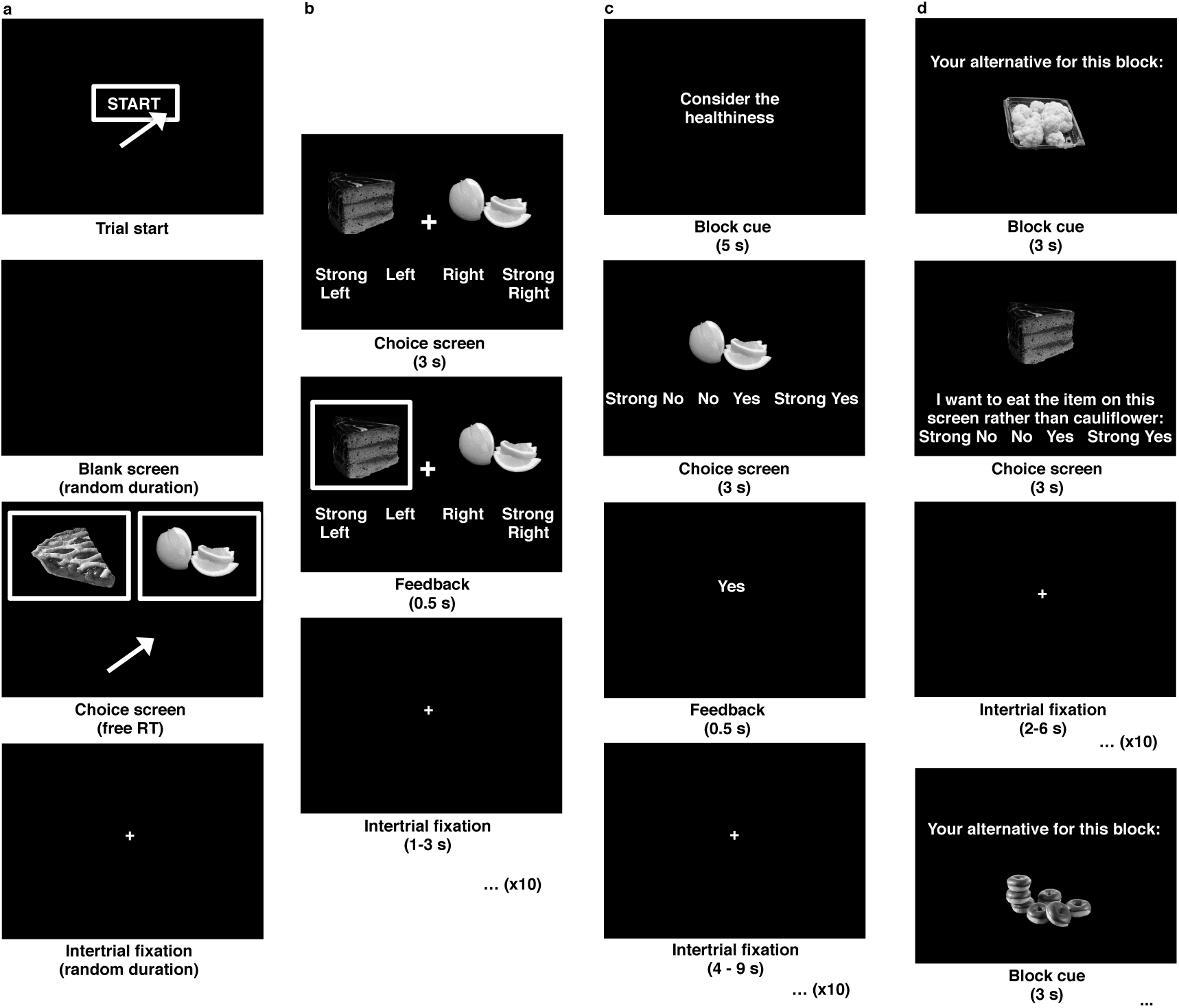
Details of the different food choice tasks used in each study. **(a)** On each trial in the mouse response trajectory (MRT) task used by Sullivan et al. ^16^, participants first saw a start screen and had to respond by continuously moving the mouse toward the option they wanted to choose until they reached the box that contained the desired item. **(b)** In our gambles plus food choices (GFC) study, participants chose between two foods without ever being instructed to think about healthiness. They had up to 3 seconds to make their choice on a 4-point scale ranging from strongly prefer left to strongly prefer right. Intermixed between the food choices were trials on which participants had to select between decks of cards for monetary rewards. **(c)** In the instructed attention cues (IAC) task used by Hare et al. ^18^, cues to consider a specific attribute or choose naturally were depicted for 5 seconds before each choice block of 10 trials. Participants then had 3 seconds to make their choice on a 4-point scale from strong no to strong yes. **(d)** In our TDCS study, the reference food for the upcoming block was shown for 3 seconds before each block began. During each block, a series of 10 different foods were shown together with a 4-point scale from strong no to strong yes (in favour of eating the item shown over the reference). The identity of the reference food was written in text below each alternative shown on the screen as depicted in panel **d**.

## Results

We adapted the traditional drift diffusion modelling (DDM) framework ^19–21^ to allow for each attribute in a multi-attribute decision problem to enter into the evidence accumulation process at separate times (Figure 2a). We chose the DDM as a starting point because this type of sequential sampling model is relatively simple, yet has often been shown to be useful in explaining behaviour across many domains ^21^ (see Supplementary Discussion). This modified model is a time-varying DDM (tDDM) because the separate consideration onset times for each attribute cause the drift rate to vary over time within a choice. Briefly, we add a free parameter (relative start time (RST)) estimating how quickly one attribute begins to influence the rate of evidence accumulation relative to another.

**Figure 2.**
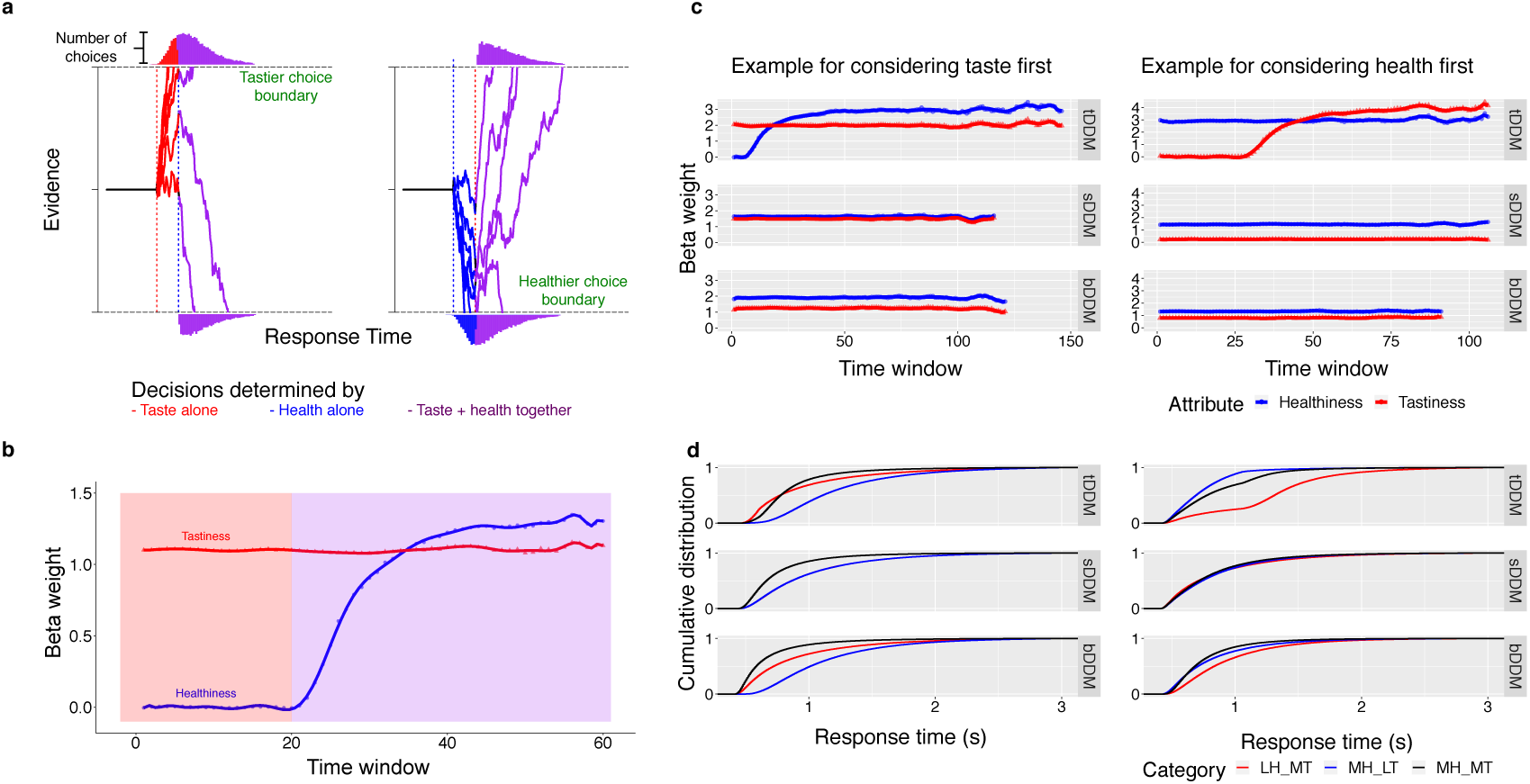
Patterns of behaviour predicted by this time-varying DDM. (**a**) The trajectories that begin as red are simulations from an agent that considers tastiness alone first before beginning to consider healthiness attributes. Those that begin as blue are for an agent that considers healthiness before taste). Our time-varying DDM includes an additional parameter, the relative start time (RST), that represents the average difference in consideration start times between attributes. If a choice boundary is not reached within the RST, then both attributes influence the decision. This is indicated by the lines becoming purple. The histograms at each boundary show the distribution of response times for each choice outcome. Note that choices in favour of the earlier-considered attribute are more frequent and faster, on average. (**b**) The y-axis shows the β_1_ (red triangles and line) and β_2_ (blue circles and line) coefficients from the logistic regression, Choice ∼ β**_0_** + β**_1_**\*Taste_difference + β**_2_**\*Health_difference + *e*, as a function of response time (RT) from a simulated agent that considers tastiness first. The x-axis represents overlapping 100 ms RT windows that slide from minimum to maximum RT in steps of 20 ms. Red shading indicates the period during which only tastiness influences the decision, and purple shading indicates choices made after both attributes are considered. (**c**) These plots are analogous to (b), but are based on simulations from the best-fitting tDDM, standard DDM (sDDM), and tastier starting-point-bias DDM (bDDM) parameters for two example participants. (**d**) The plots show the cumulative density functions of the RTs from the simulated choices in (c) as a function of choice outcome. Choice outcome abbreviations: LH_MT = less healthy but more tasty (i.e. health challenge failure); MH_LT = more healthy but less tasty (i.e. health challenge success); MH_MT = more healthy and less tasty (i.e. no challenge).

The drift rate determining the evidence update at each time step (dt = 8 ms) if taste enters first is

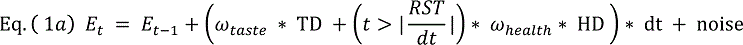

While if healthiness enters first it is

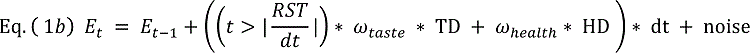

Where *E* is decision evidence, *t* is the timestep, RST = relative starting time in ms, dt = 8 ms, TD = tastiness difference, HD = healthiness difference, and ω_taste_ and ω_health_, are subjective weights for taste and health, respectively.

Thus, the times at which the weighted value differences in tastiness and healthiness attributes begin to influence the evidence accumulation rate are determined by RST. When the conditional statement 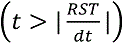 is false, it equals 0, while if true it equals 1. Multiplying one of the two weighted attribute values by 0 until 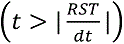 is true means that this attribute does not factor into the evidence accumulation process for the initial time period determined by |RST|. The RST parameter is defined as the consideration start time for healthiness minus the starting time for tastiness. Note that the standard, synchronous onset DDM is equivalent to the specific case of *RST* = 0.

We found that the attribute timing asynchrony estimated from response times by our tDDM was significantly associated with the results from Sullivan et al.’s mouse response trajectory analysis. Participants in that study made choices by moving a computer mouse from the bottom centre to the upper left or right corners of the screen to indicate their choices. Sullivan et al. analysed the response trajectories to determine the relative times at which health and taste attributes enter into the decision. We compared their estimates with those we computed using the tDDM for the same data (see Table 1). The time at which healthiness attributes entered into consideration was significantly correlated across the two analysis methods (r = 0.503, Posterior Probability of the correlation being positive (PP(r > 0)) = 0.991, 95% Highest Density Interval (HDI) = [0.157; 0.811], Bayes factor (BF) = 7.86), establishing face validity for the tDDM estimates.

**Table 1.**
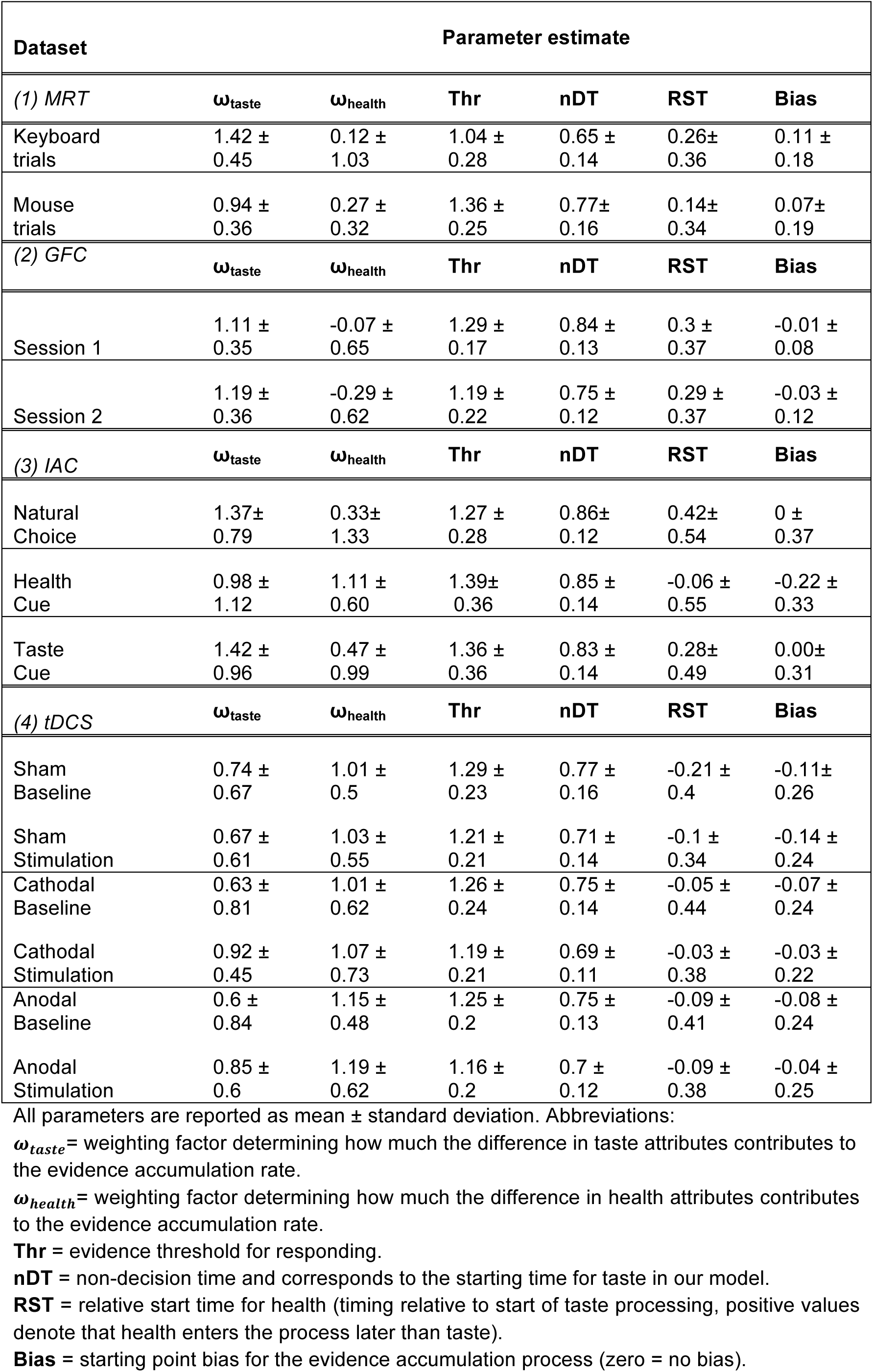
Fitted separate attribute onset tDDM parameters by study and condition

In total, we tested the tDDM in 272 participants across four datasets with different experimental conditions (mouse response trajectory choices (MRT), standard binary choices in a combined gambling and food choice (GFC) task that was repeated two weeks apart, choices following instructed attention cues (IAC) toward taste or healthiness, and choices under transcranial direct current stimulation (TDCS) (see Figure 1). The tDDM yielded a better fit to choices and response time (RT) distributions than the standard formulation of a DDM with a single, synchronous onset time (overall tDDM BIC = 280632, overall standard DDM BIC = 281909; Figure 3, Supplementary Table 1).

**Figure 3.**
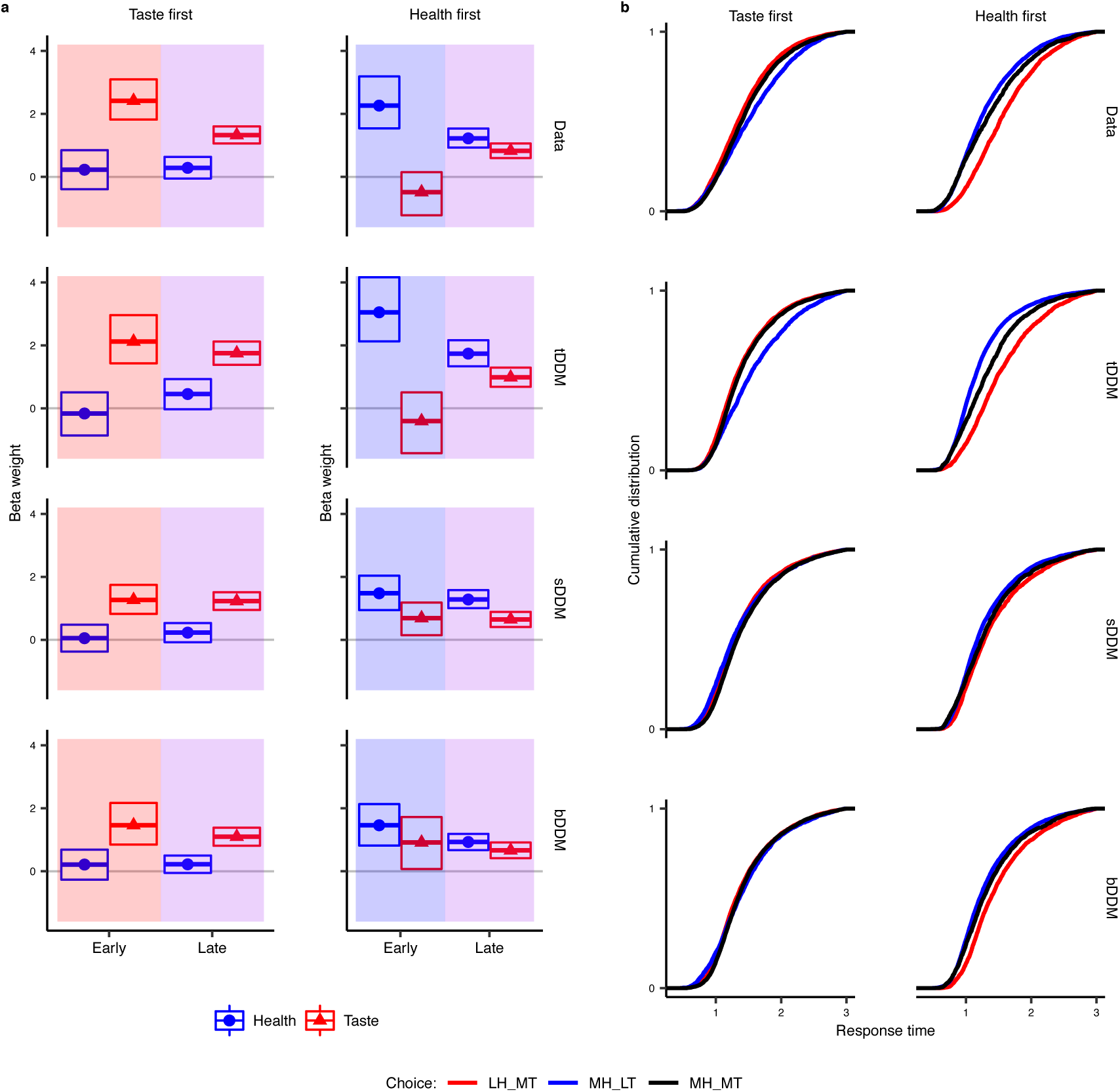
Influences of taste and healthiness attributes on choice outcomes and response times. Row 1 shows results from the empirical data and rows 2-4 show results from data simulated using the best-fitting time-varying (tDDM), standard (sDDM), and taste-starting-point-bias (bDDM) drift diffusion model parameters. The tDDM reproduces the time-dependent attribute influence and response time (RT) patterns found in the data better than either alternative model. **a**) These plots show the mean (filled shapes) and 95% highest density intervals (open rectangles) of beta weights from the logistic regressions in Supplementary Table 5 and represent the estimated influences of tastiness (red triangle) and healthiness (blue circle) attributes on choice outcomes. These influences are shown separately for early (< 1s) and later (> 1s) response times and participants that are estimated to consider taste or health first. The blue, red, and purple background shading corresponds to Figure 2 and indicates periods where, on average, health alone, taste alone, or both taste and health are expected to have a significant influence on choices. **b**) These plots show the cumulative distributions of response times (RT) observed in the empirical and simulated data. Once again, the first and second columns show data from participants who are estimated to consider taste or healthiness first, respectively. The three colours indicate choice outcomes in favor of 1) less healthy but more tasty (LH_MT) in red, 2) more healthy but less tasty (MH_LT) in blue, or 3) both more healthy and more tasty (MH_MT) in black. Choices in favour of the option rated as less healthy and less tasty were rarely made (less than 5% of trials) and are omitted for clarity.

Parameter recovery tests demonstrated that choice and RT patterns simulated using known values of the standard and tDDM could be recovered in each case (see Extended Data Figure 1, Supplementary Figure 1, and Supplementary Results). In other words, our estimation procedures for the tDDM yield accurate parameter estimates. Critically, the parameter recovery tests also showed that earlier (later) onset of evidence accumulation can be distinguished from stronger (weaker) weighting of evidence (Extended Data Figure 1e). Furthermore, the tDDM with separate onset times also generated significantly better out-of-sample predictions for food choices than the standard DDM. The mean squared error for this tDDM (0.163) was lower than that of the standard DDM (0.170) (posterior probability of greater accuracy for this tDDM versus a standard DDM = 0.97; see also Supplementary Table 2). Thus, fitting the tDDM allows for more accurate predictions about out-of-sample dietary choices.

Adding the separate onset time feature allows the model to capture important choice and RT patterns. Specifically, different onset times for the two attributes can explain the fact that the relative contribution of tastiness and healthiness attributes to the evidence in favour of one food changes during the decision process. This change in the relative weighting of taste versus healthiness in our data can be seen in the simulations depicted in Figure 2b-d, and when computing a logistic regression model that estimates the influence of taste and healthiness on participants’ choices made before or after both attributes were estimated to have begun being considered on average (Figure 3, Supplementary Tables 3 and 4). Shared onset time DDMs cannot replicate the effect as shown in Figures 2, 3, Extended Data Figure 2, and Supplementary Table 5. We note that separate attribute consideration onset timing is a general feature that could be added to many other types of sequential sampling models in addition to the DDM, e.g. ^12, 22–28^.

**Table 3.** Effects of tDCS over left dlPFC on tDDM parameters.

|  | Mean difference | 95% HDI | Posterior probability | Bayes factor |
| --- | --- | --- | --- | --- |
| **A. Taste weighting ($\omega_{\text{taste}}$)** |  |  |  |  |
| Baseline - Anodal tDCS | -0.094 | [-0.23 0.043] | 0.081 | 0.047 |
| **Cathodal tDCS - Baseline** | **0.138** | **[0.027 0.248]** | **0.993** | **5.748** |
| Baseline - Sham tDCS | 0.072 | [-0.087 0.23] | 0.815 | 0.33 |
| ΔSham - $\Delta$ Anodal | -0.185 | [-0.41 0.047] | 0.053 | 0.091 |
| **$\Delta$ Cathodal - $\Delta$ Sham** | **0.215** | **[0.014 0.42]** | **0.982** | **8.333** |
| **B. Health weighting ($\omega_{\text{health}}$)** |  |  |  |  |
| Anodal tDCS - Baseline | 0.098 | [-0.023 0.221] | 0.941 | 0.267 |
| Baseline - Cathodal tDCS | -0.074 | [-0.246 0.102] | 0.197 | 0.094 |
| Sham tDCS - Baseline | 0.025 | [-0.082 0.13] | 0.685 | 0.188 |
| $\Delta$ Anodal - $\Delta$ Sham | 0.063 | [-0.093 0.223] | 0.787 | 0.261 |
| $\Delta$ Sham - $\Delta$ Cathodal | -0.039 | [-0.241 0.164] | 0.349 | 0.149 |
| **C. Relative start time of health (RST)** |  |  |  |  |
| Baseline - Anodal tDCS | -0.002 | [-0.098 0.093] | 0.484 | 0.144 |
| Cathodal tDCS - Baseline | 0.021 | [-0.095 0.135] | 0.648 | 0.193 |
| Baseline - Sham tDCS | -0.103 | [-0.225 0.018] | 0.047 | 0.056 |
| ΔSham - $\Delta$ Anodal | 0.098 | [-0.057 0.251] | 0.895 | 0.764 |
| ΔCathodal - $\Delta$ Sham | -0.081 | [-0.25 0.086] | 0.171 | 0.106 |

This table reports changes in the separate attribute consideration onset tDDM relative starting times (RST) or weighting parameters ( $\omega_{\text{taste}}$ , $\omega_{\text{health}}$ ) as a result of tDCS over the left dIPFC. Rows in bold indicate changes that are significantly different from zero. The $\Delta$ symbol always indicates a difference score equal to the value in the stimulation minus the baseline session within a given condition. Rows containing this symbol report differences of differences across conditions. Mean differences (or differences of differences) and their 95% highest density intervals (HDI) were computed based on 100,000 samples drawn from the posterior distributions of each parameter. The third column displays the posterior probabilities that differences are greater than zero. All comparisons were made so that *a priori* predicted effects would be positive.

This feature of our tDDM differs in important ways from other types of multi-process sequential sampling models that include a combination of fast automatic processes and slower deliberate processes (e.g. dual process, fast guess, Ulrich diffusion model for conflict tasks (DMC)) ^29–32^. These other frameworks can account for changes in the way evidence is accumulated over time in certain cognitive tasks, but are fundamentally inconsistent with our food choice data. First, responses made before the second attribute enters into consideration are similarly or even more sensitive to the level of the first attribute, relative to choices made after both attributes begin to be considered. This indicates that these choices are not random guesses or prepotent or habitual responses. Second, the data from the instructed attention cue experiment described below show that whether tastiness or healthiness is considered first is not automatic. Thus, modifying sequential sampling models to allow different attributes to enter into a deliberate consideration process at separate times is more appropriate to explain the outcomes and response times from the goal-directed choice process studied here.

In the paragraphs above, we established the face validity (i.e. correspondence to the mouse trajectory analysis), accuracy (i.e. good parameter recovery), and predictive utility (i.e. improved out-of-sample predictive accuracy relative to the standard DDM) of our modelling approach. Next, we used a tDDM to test several fundamental questions about how attribute timing and weighting work together, or potentially separately, to influence choice outcomes during healthy choice challenges.

### Are more abstract attributes considered later in the choice process?

One may assume that for dietary choices, the relative start time of the more abstract attribute (healthiness) will lag behind the more concrete and immediately gratifying attribute of taste. However, our results indicate that this is not the case. Pooling the data across all studies, we found that the posterior probability that healthiness entered into consideration later than tastiness was only 0.48 (mean difference in starting times = 0.001 seconds; 95% HDI = [−0.05; 0.06; BF for RST > 0 = 0.21). In total, only 130 out of 272 participants (48 %) had relative-start-times for healthiness attributes that were delayed relative to those for tastiness (Extended Data Figure 3). We used the linear regression model in equation (3) to test the relationship between RST and other tDDM parameters. The relative-start-time parameter was related to both the tastiness and healthiness weights as well as to the starting point bias parameter (Supplementary Tables 6 and 7), but overall the linear combination of other tDDM parameters explained only 30% of the variability in relative start times across participants.

### Are individual relative start times and attribute weights stable over time?

We tested whether tDDM parameters provide good estimates of stable individual characteristics by comparing parameters estimated from food choices made by the same participants two weeks apart. Previous work has shown that the test-retest reliability of choice outcomes in the food choice task is high when participants repeat the same incentivised choices a few days or one month apart ^33^. In contrast, within each session of our GFC study, participants (N = 37) faced 150 trials consisting of a choice between two randomly paired food items. In other words, participants did not complete the exact same set of trials on the two visits, but instead the food pairings varied randomly. This precludes a direct comparison of choice outcomes in the two sessions. Nevertheless, individual characteristics inferred from the tDDM parameter fits were quite consistent over time. The relative weighting of taste and healthiness attributes (i.e. taste > health or vice versa) was the same for 92% of participants across both visits, while the attribute considered first (i.e. RST) was consistent in 76% of participants. Furthermore, tDDM parameters fit to choices in session 1 accurately predicted new food choices made in session 2 two weeks later 77% of the time. For comparison, in-sample predictions for session 2 choices based on tDDM parameters fit to those same choices were correct 78% of the time. These results indicate that taste versus health weighting and consideration onset times may be relatively stable individual characteristics, at least in the absence of experimental manipulations or interventions designed to alter these choice processes.

### Effects of attention cues on attribute weights and relative-start-times

Next, we examined whether directing attention toward either healthiness or tastiness could change the time at which those attributes enter the decision process and if changes in timing were linked to changes in weighting strength. This analysis was motivated by previous findings ^18^ that directing attention to the healthiness aspects of a food item resulted in substantial changes in choice patterns (Figure 4a). In this instructed attentional cues (IAC) experiment, instructive cues highlighted health, taste, or neither attribute for explicit consideration during the upcoming block of 10 food choices. We refer to these three block types as health-cued (HC), taste-cued (TC), and natural-cued (NC). The original analysis of these choice data focused on the regression weights for taste and health attributes in each choice condition but did not consider that the cues might change the relative times at which these attributes entered into the choice process. Our goal was to determine how potential alterations in attribute timing and weighting contributed to the observed changes in choice behaviour during health cue relative to natural blocks.

**Figure 4.**
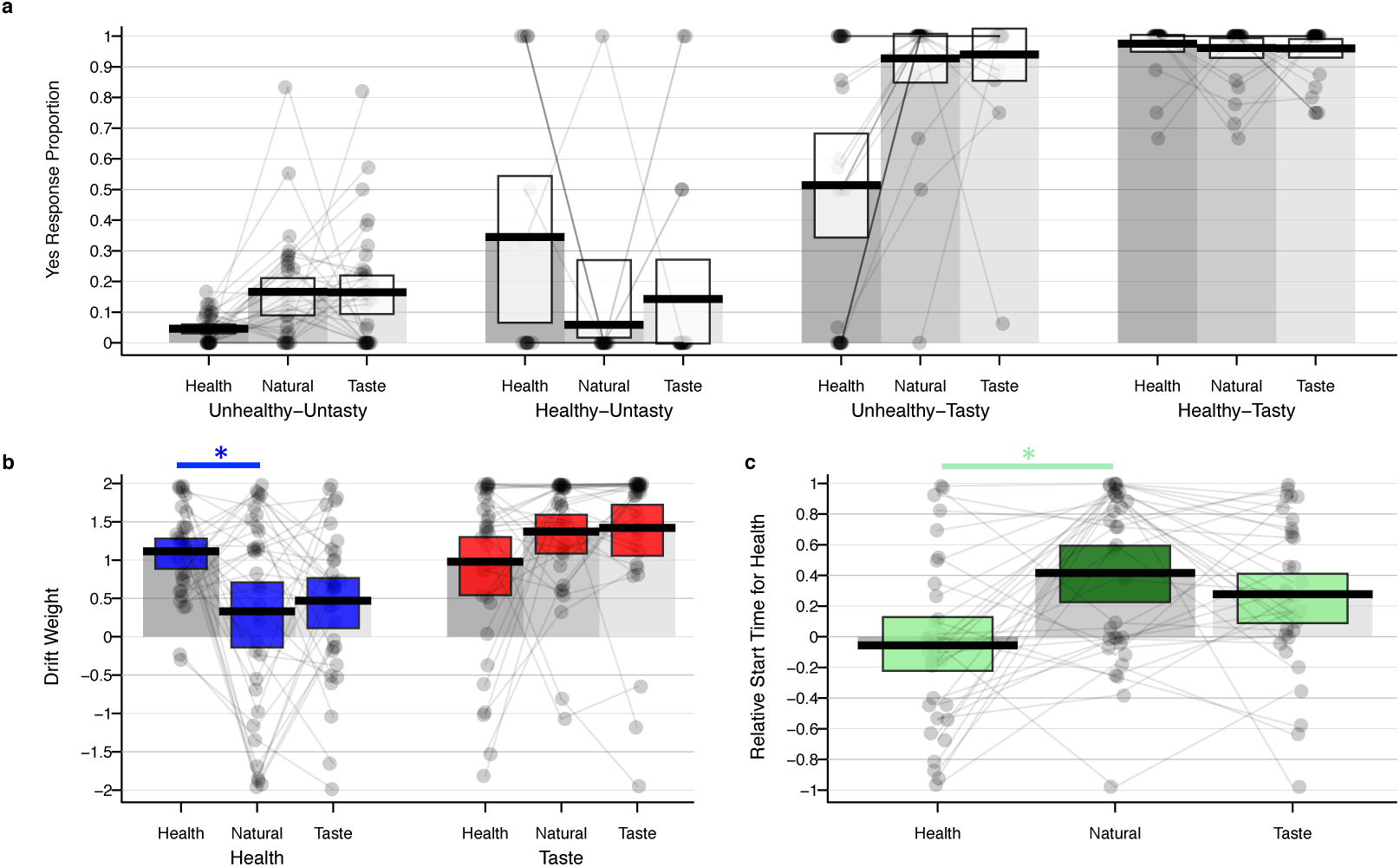
Choice patterns and separate attribute consideration onset tDDM parameter estimates for the IAC study (n=33 participants) by condition. **(a)** Proportion of times that subjects chose to eat the food (i.e., they responded “yes” or “strong yes”) as a function of attention cue type (Health, Natural, or Taste) and taste-health combination of food under consideration (Tasty or Untasty crossed with Healthy or Unhealthy). In terms of mean choice proportions, directing attention towards healthiness decreased the proportion of choosing healthy-untasty items and decreased the proportion of choosing unhealthy-tasty items compared to the natural condition. The changes in choices during Health blocks were accompanied by higher weights and faster relative-start-times for healthiness. **(b)** Compared to the natural condition, attention cues to health resulted in a higher relative drift weight (arbitrary units) for the corresponding health attribute compared to the natural condition (blue shading). There was no significant change in the relative drift weights of the two attributes during the taste cue blocks (red-shading). **(c)** Attention cues to health also led to a faster relative start time (seconds) for health attributes compared to natural blocks (green shading). Once again, there was no significant difference in relative timing between natural and taste blocks. For all plots, the dots within each column represent the value for a single participant in the sample. Darker shading indicates that multiple participants share the same value for that parameter. Black horizontal bars indicate condition means and white, blue, red, or green shaded rectangles indicate the 95% HDIs for each measure. The grey shaded bars in each plot serve to visually separate the columns for each condition and highlight the zero-points on the y-axes.

First, we found that attention cues changed both the relative weighting and timing of taste and healthiness attributes. Compared to the natural choice blocks, 70% of participants reversed their relative weighting of taste and healthiness in taste or health cue blocks (i.e., went from taste > healthiness to taste < healthiness weight or vice versa), and 64% switched whether they considered tastiness or healthiness first. There was no significant difference in how often participants reversed the order of relative weights (switching from ω_taste_ > ω_health_ to ω_taste_ < ω_health_, or vice versa) compared to how often they reversed the order of relative onset times (switching from taste first to healthiness first, or vice versa) when moving from attribute-cued and natural-choice blocks (PP(weight reversal more prevalent than timing reversal) = 0.70; BF = 1.4).

Focusing our analyses on the health cue blocks that showed a significant change in choice outcomes compared to natural blocks (Figure 4a), we found that, on average, cuing attention to health attributes both significantly increased the magnitude of participants’ weights for healthiness and sped up the time at which health entered into the accumulation process (relative to taste, i.e., relative start times) (Figure 4b-c; Table 2). These results demonstrate that both the timing and weighting of taste and healthiness attributes can be flexibly and rapidly changed in response to the attention cues preceding every block of 10 choices.

**Table 2.** Changes in separate attribute consideration onset tDDM parameters between attention cued conditions.

|  | Mean difference | 95% HDI | Posterior probability | Bayes factor |
| --- | --- | --- | --- | --- |
| *A. Taste weighting ($\omega_{taste}$)* |  |  |  |  |
| Natural - Health | 0.354 | [-0.113 0.832] | 0.933 | 1.251 |
| Taste - Health | 0.455 | [-0.088 1.003] | 0.951 | 1.122 |
| *B. Health weighting ($\omega_{health}$)* |  |  |  |  |
| **Health - Natural** | **0.746** | **[0.188 1.325]** | **0.995** | **12.818** |
| **Health - Taste** | **0.633** | **[0.245 1.028]** | **0.999** | **39.041** |
| *C. Relative start time of health (RST)* |  |  |  |  |
| **Natural - Health** | **0.469** | **[0.2 0.748]** | **0.999** | **60.868** |
| **Taste - Health** | **0.336** | **[0.121 0.548]** | **0.999** | **26.655** |

This table shows the effects of attention cues on the tDDM parameters estimated from choice data in the IAC study. Changes in relative starting times (RST) or weighting parameters ( $\omega_{taste}$ , $\omega_{health}$ ) induced by the experimental conditions that are shown in bold were significantly different from zero. Mean differences and their 95% highest density intervals (HDI) were computed based on 100,000 samples drawn from the posterior distributions of each parameter<sup>34</sup>. The third column displays the posterior probabilities that differences are greater than zero. All comparisons were made so that *a priori* predicted effects would be positive. Abbreviations: $\omega_{taste}$ = weighting factor determining how much the difference in taste attributes contributes to the evidence accumulation rate. $\omega_{health}$ = weighting factor determining how much the difference in health attributes contributes to the evidence accumulation rate.

### Dissociating attribute weighting strengths and timing at the neural level

We next addressed the question of whether attribute weighting strength and timing are implemented by dissociable neural processes. We did so by analysing data from an experiment applying cathodal, anodal, or sham transcranial direct current stimulation (tDCS) over left dlPFC during food choices (see Methods section for details). Numerous neuroimaging and electrophysiological studies have reported correlational evidence for a role of the dlPFC in multi-attribute choice ^35–38^. There is also ample evidence that applying brain stimulation (both transcranial direct current and magnetic) over multiple different sub-regions of the left or right dlPFC is associated with changes in several different forms of multi-attribute decision making ^39–45^. Here, we applied tDCS over a region of the left dlPFC that has been shown to correlate with individual differences in health challenge success rates and multi-attribute decisions more generally ^18, 46–54^ in order to uncover the mechanistic changes in the choice process caused by tDCS over this particular region. Stimulation over this region of left dlPFC did not significantly change measures of working memory, response inhibition, or monetary temporal discounting in our participants (see Supplementary Results).

Previous studies suggest that the effects of stimulation over left dlPFC are strongest on trials in which the participant does not strongly favour one outcome over the other (i.e., stimulation effects are greatest in difficult choices) and depend on baseline preferences over the rewards ^40, 44^.

Therefore, we restricted our analysis of health challenge success to trials in which the predicted probability of choosing the healthier food was between 0.2 and 0.8 and focused on the difference in behaviour between baseline and active-stimulation choice sessions. Specifically, we computed a Bayesian hierarchical logistic regression model that accounted for both stimulation type and the healthiness and tastiness differences on each trial in the tDCS dataset (see equation (4) in the Methods for details). We compared the interaction effects measuring changes in each participant’s health challenge success from the pre-stimulation baseline to the active stimulation condition for cathodal and anodal versus sham simulation groups. This revealed a greater decrease in health challenge success under cathodal relative to sham stimulation (Supplementary Table 8; regression coef. = −0.32 ± 0.15, 95% HDI = [−0.58; −0.08], PP(cathodal polarity X active stimulation interaction coef. < 0) = 0.98), but no change in health challenge success for anodal relative to sham stimulation (regression coef. = −0.03 ± 0.15, 95% HDI = [−0.28; 0.22], PP(anodal polarity X active stimulation interaction coef. > 0) = 0.4). There was also a main effect within the cathodal stimulation group indicating that these individuals had fewer health challenge successes when making food choices under active stimulation compared to their pre-stimulation baseline choices (regression coef. = −0.31 ± 0.15, 95% HDI = [−0.55; −0.08], PP(active stimulation < 0) = 0.9999). Thus, we find that inhibitory stimulation over left dlPFC leads to fewer health challenge successes (see Figure 5a as well).

**Figure 5.**
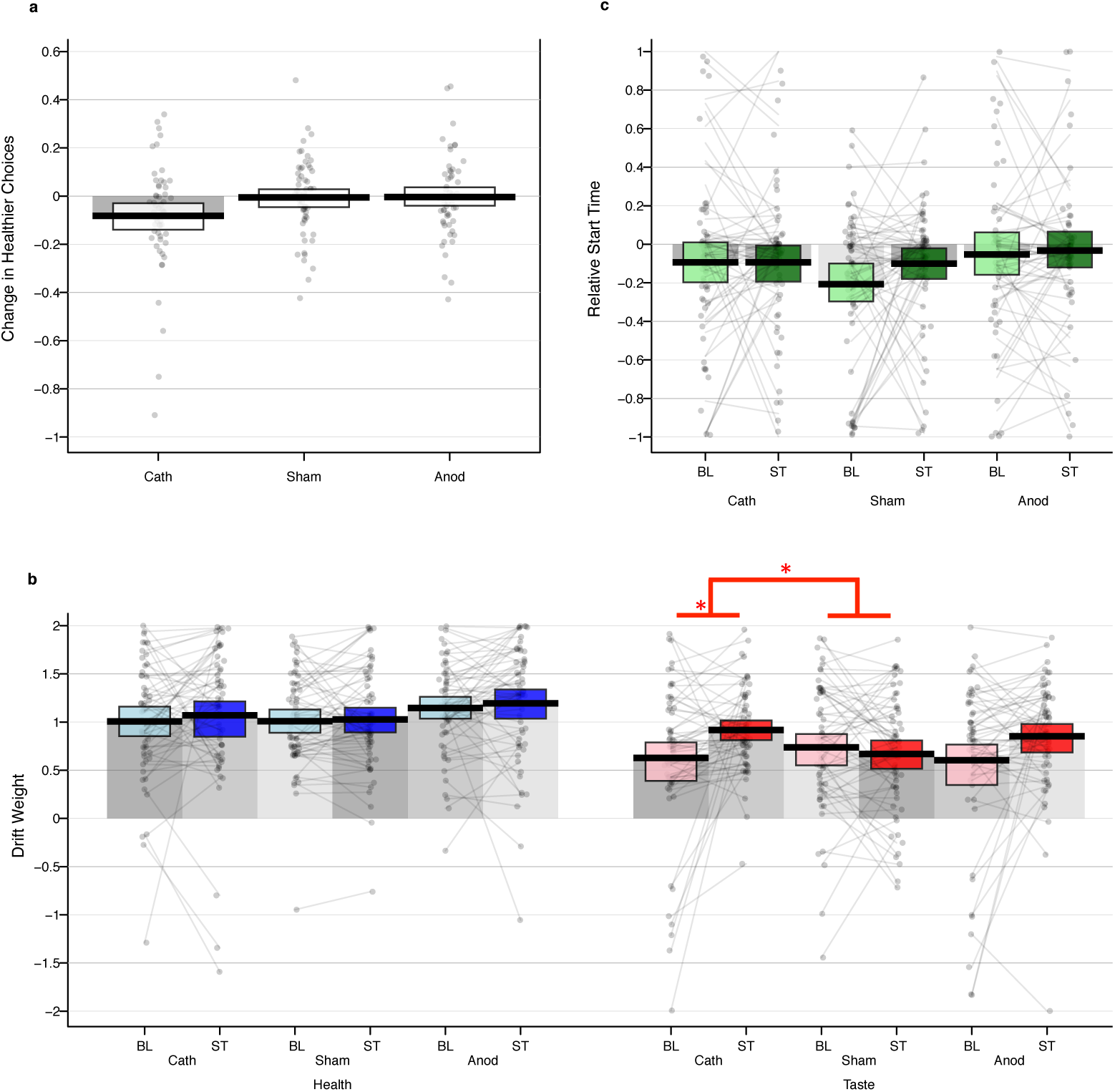
Changes in health challenge success and tDDM parameter estimates following tDCS over left dlPFC. **(a)** This plot shows the raw changes in health challenge success under stimulation compared to baseline across stimulation groups (n=58 anodal, n=57 cathodal, n=59 sham). Unlike the regression summarized in Supplementary Table 8, the effects shown in this plot do not account for the taste or health differences in each choice. Each dot represents the difference between active stimulation or sham and baseline in one participant. Left dlPFC-targeted cathodal stimulation significantly decreased health challenge success (mean decrease = 5.7% ± 0.02%, 95% HDI = [−10%; −1.5%], PP(active cathodal stimulation < 0) = 0.995, Bayes factor = 6.41). Note that we tested the change under cathodal stimulation using a method ^34^ that is robust to outliers such as the two extreme participants near −1. There were no significant differences in healthy choices under anodal or sham stimulation. **(b)** Cathodal stimulation increased the weighting of taste attributes (ω_taste,_ red shading on right) relative to baseline choices, PP(Cath ST > Cath BL) = 0.99, Bayes factor = 5.75). This change from baseline was greater under cathodal stimulation than sham, PP((Cath ST – Cath BL) > (Sham ST – Sham BL)) = 0.98, Bayes factor = 4.20). The red lines and stars highlight this main effect and interaction. Anodal stimulation did not lead to significant changes in attribute weighting parameters, and neither tDCS protocol affected drifts weights for healthiness (ω_health_, blue shading on left). The weighting strength parameters are plotted in arbitrary units. **(c)** tDCS had no significant effect on the RST parameters (plotted in seconds, green shading). Black horizontal bars indicate group means and rectangles indicate the 95% HDIs for each effect. The grey shaded bars in each plot serve to visually separate the columns for each condition and highlight the zero-points on the y-axes.

In order to elucidate the changes in choice processes caused by the stimulation, we fit the separate attribute consideration onset tDDM to dietary choices made during the pre-stimulation baseline and active or sham tDCS sessions. When testing how the tDDM parameters changed between baseline and active stimulation sessions in each group, we found that the cathodal group had increased weighting of taste attributes under stimulation compared to baseline choices (mean difference = 0.14, HDI = [0.03; 0.25], PP(Cath active > Cath baseline) = 0.99, BF= 5.75; Figure 5a) and that the change from baseline was greater under cathodal stimulation than sham (mean difference = 0.21, 95% HDI = [0.01; 0.42], PP(Δ Cath > Δ Sham) = 0.98, BF = 4.20).

Crucially, the relative-start-time parameters were unaffected during left dlPFC-targeted cathodal tDCS (Table 3, Figure 5b). Moreover, the tDCS-induced changes in taste relative to health weighting parameters and relative start times were not significantly correlated (r = −0.07, 95% HDI = [−0.325; 0.188], PP(r > 0) = 0.30, BF = 0.210). Consistent with the lack of significant change in choice behaviour under anodal tDCS, we found no significant changes in any tDDM parameter under anodal stimulation (Table 3). In summary, we found that cathodal tDCS over left dlPFC changed the relative decision weight placed on taste attributes, but not the speed with which taste, relative to healthiness, began to influence the choice process (Table 3).

## Discussion

We have shown that separable mechanisms determine the degree to which an attribute affects the evidence accumulation rate (weighting strength) and the relative speed with which it begins to do so (timing). Measuring each of these distinct processes helps to explain individual differences in dietary choices at baseline as well as how behavioural and neurophysiological manipulations effect changes in the decision process. Thus, both attribute timing and weighting strength must be examined if we seek to better understand decision making at the mechanistic level.

The clearest evidence that timing and weighting strength are dissociable comes from our tDCS experiment showing that stimulation over the left dlPFC caused a change in the weights placed on the taste factor, but not the timing of taste versus healthiness attributes during dietary choices.

Moreover, changes in the relative weighting and the relative timing of each attribute between baseline and cathodal stimulation sessions were not significantly correlated, further indicating that the neural mechanisms altered by our tDCS protocol were specifically related to attribute weighting (further implications of these results are included in the Supplementary Discussion).

In our current work, for example, we found that the relative importance given to a specific attribute, as well as its speed in entering into the choice process, could be altered by instructions that directed attention to that attribute. Although a large body of work has established that value construction and comparison processes are malleable and subject to attention, perceptual constraints, and other contextual factors ^9–11, 57, 58^, the influence of attribute consideration timing within a given decision is rarely discussed or directly tested. Query theory ^59, 60^ is a notable exception in that it explicitly posits that the order in which attribute values are queried from memory or external sources will bias value construction and choice processes because the recall of initial attributes reduces the accessibility of subsequent attributes. Although the current data cannot be used to address the question directly, future experiments may address the important mechanistic question of whether or not memory retrieval is a driving factor in the consideration onset asynchronies revealed by the separate attribute consideration onset tDDM.

Despite open questions about the relationship between memory and relative starting times, our finding that attribute consideration start times are asynchronous lends strong support to the idea that choices are made based on comparisons of both separate attribute values as well as overall option values. Hunt and colleagues ^11^ demonstrated that a hierarchical sequential sampling process that operates over both separate attribute and overall option values explains risky choice behaviour and brain activity better than models operating only on integrated values. Reeck and colleagues ^10^ have shown that individual variation in temporal discounting can be explained by patterns of information acquisition that support attribute-wise or option-wise comparisons; moreover, their study shows that choices can be made more patient by an experimental manipulation that promotes attribute-wise comparisons compared to one promoting option-wise comparisons. Together, these results and others (e.g. Roe, et al. ^27^ and Bhatia ^58^) indicate that attribute-level comparisons play an important role in determining choice outcomes.

Hierarchical attribute and option-level comparisons are implicit in our specification of the separate attribute consideration onset tDDM because the choice outcome and response time are determined by a weighted sum of the differences in attribute values. However, we show that attribute-level comparisons do not all begin at the same point in time, and that the magnitude of the difference in relative start times across attributes influences option-level comparisons and choice outcomes.

Our results raise important questions about how attribute weighting strengths and onset timing jointly influence choice outcomes: How should we interpret choices in which the outcome is determined by the advantage in relative timing as opposed to weighted evidence? Could this be strategic use of cognitive flexibility to align decision making with current goals or should we consider such outcomes to be mistakes? Traditionally, a weighted combination of all attribute values is assumed to yield the “correct” choice ^61^.

If the weighting strength on each attribute is appropriate, then any asynchrony in onset timing could produce suboptimal choices (i.e., choices in favour of options with a lower weighted sum over all attribute values than another available alternative). In that sense, it is surprising that we find substantial attribute onset asynchrony in healthy young adults and that, in individuals striving to maintain a healthy lifestyle (i.e., the sample recruited for our tDCS experiment), a higher level of asynchrony is associated with better health challenge success. However, this view is predicated on the assumption that the attribute weighting strengths are appropriate for the current goal or context.

On the other hand, it is possible that shifts in the timing of attribute consideration can be used to achieve the desired outcome. Suppose that a decision maker knows (not necessarily explicitly) that her standard attribute weights are inconsistent with her current decision context or goal, and adjusting those weights by the necessary amount is costly or unlikely. In that case, shifting the relative onset timing could be an effective means of reducing effort and improving the chances of making a goal-consistent choice. In other words, altering the relative starting times may be a form of proactive control ^62, 63^. For example, a decision maker who goes on a diet may find it difficult to convince herself that she does not like the taste of ice cream and/or to constantly trade off this delicious taste against the downsides of excess sugar and fat. An alternative way to bring about health challenge success in this situation may be to adjust the process(es) that determine relative start times for healthiness and tastiness, to focus on the healthiness of each alternative option alone for a brief period in order to forgo extremely unhealthy options (without putting in time or effort to compare taste benefits to health costs).

The use of timing differences as described above would be consistent with at least two existing theories on the role of attention in cognition. First, it is consistent with the idea that rational inattention strategies ^64–66^ can be employed as a means of reducing effort costs. Specifically, if the time advantage for healthiness is large enough, then one could theoretically decide against eating an unhealthy food before even considering its tastiness and thus not experience temptation or conflict. Second, the idea that distinct processes determine consideration onset times and weights for different attributes is paralleled in theories of emotion and food-craving regulation that posit separate attention deployment and stimulus appraisal steps (e.g. ^36, 67, 68^).

However, we don’t yet know if strategic use of attribute consideration onsets or related processes actually happen, nor if adjusting the process determining relative onset times is, in fact, less effortful or more likely to succeed than strategies that attempt to alter the attribute weighting strengths.

Altering the processes that determine the relative onset times could be a means or a result of delaying and reducing attention. However, although we found that both cueing attention to healthiness and having the goal of maintaining a healthy lifestyle (tDCS sample vs. all others) were associated with faster average onset times for healthiness attributes, we don’t know yet if relative onset times can be manipulated as part of a deliberate strategy. It is also important to note that the response to healthiness cues was heterogeneous in the sense that, although most participants made healthy choices more often following those cues, some participants changed only attribute weights or only attribute start times in favour of healthy choices, rather than both. Further research is needed to understand why individuals responded to these cues in different ways.

The ability to understand or predict how an intervention or policy change will affect choice processes and their outcomes for specific individuals or groups of people is important for any program hoping to promote behavioural change, for example in domains such as health, crime, or financial stability. Greater knowledge of the cognitive and neural mechanisms that drive choices in specific individuals is an important step toward this understanding ^69^. Our findings demonstrate that when a specific attribute begins to influence the decision process - a factor that has been generally neglected - is an important determinant of choice outcomes. They also suggest that examining relative differences in attribute start times may prove to be useful in understanding why interventions and policies work in some cases (e.g., for specific individuals or groups) but not in others, and may help to increase their effectiveness. Overall, the work we present here provides both a concrete advancement in our knowledge of multi-attribute choice processes and a functional set of computational modelling tools that can be applied to extract deeper mechanistic insights from data on choice outcomes and response times.

## Methods

For all data sets in which we relied on published studies, we included the final reported sample in our analyses. For these studies, we will describe the methodological details relevant for our analyses and refer the reader to the published papers for any further details. All participants provided written informed consent in accordance with the procedures of the Institutional Review Board of the California Institute of Technology, the Institutional Review Board of the Faculty of Business, Economics and Informatics at the University of Zurich, or the Ethics Committee of the Canton of Zurich. All participants received a flat fee to compensate for their time in addition to the food they chose.

### Data set 1 – Mouse response trajectories (MRT)

We use the choice and response time data from the study of Sullivan et al. ^16^ to test the face validity of our time-varying sequential sampling model. These data are openly available at https://osf.io/jmiwn/. All participants in the MRT sample were healthy adults and had no specific dietary restrictions. Before making any choices, they were reminded of the importance of healthy eating by reading a short excerpt from WebMD.com before starting the choice task.

#### Participants

The experiment was approved by the Institutional Review Board of the California Institute of Technology. Twenty-eight (7 female) healthy adult participants completed the study.

#### Procedure

Participants were asked to fast for 4 hours prior to the study. They first rated 160 foods for taste and health on a 5-point Likert scale with values from −2 (“very little”) to +2 (“very much”). After these ratings, participants were asked to read a short text from WebMD.com on the beneficial effect of healthy eating, in order to increase the frequency with which they tried to succeed in health challenges in the following dietary choice task. In the choice paradigm, participants made 280 choices between two foods on the screen (see Figure 1a). The selection ensured that food pairs would represent all possible combinations of taste and health ratings equally. After each block of 40 choices, participants could take a short break. In 240 trials, participants used the mouse to answer, while in the remaining 40 trials, they answered with the keyboard. In mouse trials, participants had to click the “Start” box at the bottom of the screen to initiate the trial. The cursor reappeared after a random waiting period of 0.2 to 0.5 seconds. From this point on, participants had to move the mouse continuously towards the food they wanted to select. They were instructed to answer as quickly and accurately as possible. A random fixation time of 0.4 to 0.7 seconds separated the trials. In keyboard trials, participants selected food items by pressing the left or right choice keys. At the end of the study, one randomly selected trial was paid out and participants were asked to stay in the lab for 30 minutes or until they had eaten their obtained food.

### Data set 2 – Gambles plus food choices (GFC)

Data for this behavioural study (gamble plus food choice, GFC) were collected from the same individuals in two testing sessions two weeks apart. The two sessions were run on the same weekday and daytime in a two-hour visit in the afternoon. Participants in this study were healthy and did not have any specific dietary restrictions. During the study, they chose naturally and were neither reminded about eating a healthy diet nor encouraged to eat healthy in any way.

#### Participants

The Study was approved by the Institutional Review Board of the University of Zurich’s Faculty of Business, Economics and Informatics. Thirty-seven participants (17 female, mean age = 22.6 ± 3 years SD) were included in this study. A pre-screening procedure ensured that all participants regularly consumed sweets and other snack foods and were not currently following any specific diet or seeking to lose weight. All participants were healthy and had no current or recent acute illness (e.g., cold or flu) at the time of the study. All participants complied with the following rules to ensure comparability across the study sessions: They got a good night’s sleep and did not consume alcohol the evening before the study. On the study day, they took a photograph of the small meal that they consumed 3 hours before the appointment, and sent this photo to the experimenter. One day before the second study session, participants received a reminder about the rules above and were asked to consume a small meal before their second appointment that was equivalent to their meal before the first test session. Participant received 37.5 CHF (approx. 39 USD) for each session.

#### Procedure

Participants were asked to eat a small meal of approximately 400 calories 3 hours prior to their appointment and to consume nothing but water in the 2.5 hours before the study started. In the laboratory, participants first rated 180 food items for taste and health. They then made 150 food choices, one of which was randomly selected to be realised at the end of the experiment. On each trial, the screen showed 2 foods next to each other and participants chose the food they wanted to eat using a 4-point scale, picking either “strong left”, “left”, “right”, or “strong right” (Figure 1b). The pairing order and positions of the foods on the screen (left vs. right) were completely randomized, and the allocation algorithm ensured that one of the foods would be rated as healthier than the other. Participants had 3 seconds to make their choice, with a jittered interval of 1-3 seconds fixation between trials. Between blocks of dietary decisions, participants played a game in which they had to guess cards for monetary rewards. We ignore the card guessing choices for the analyses presented here. At the end of the experiment, participants stayed in the laboratory for an additional 30 minutes during which they ate the food they obtained during the study. Note that participants on the second day saw a new set of choice options that was created based on the taste and health ratings they gave on that second day, using the same allocation algorithm as in session 1.

### Data set 3 – Instructed attention cues (IAC)

In order to determine how attention cues affected attribute timing and weighting, we re-analysed data from Hare, et al. ^18^. Participants in this study were not following a specific health or dietary goal in their everyday life, but received a cue to think about the healthiness or tastiness of the foods before deciding on a subset of choices in the study.

#### Participants

The study was approved by the Institutional Review Board of the California Institute of Technology. Thirty-three participants (23 female, mean age 24.8 ± 5.1 years SD) were included. Screening ensured that they were not currently following any specific diet or seeking to lose weight. All participants were healthy, had no history of psychiatric diagnoses or neurological or metabolic illness, were not taking medication, had normal or corrected-to-normal vision, and were right-handed.

#### Procedure

Participants were instructed to fast and drink only water in the 3 hours prior to the study. In this experiment, participants made a series of 180 choices within an MRI scanner while BOLD fMRI was acquired. The experiment had three conditions with 60 trials each that were presented in blocks of 10, with the order of blocks and foods shown within blocks fully randomized for each participant. Each food was shown only once (Figure 1c).

In condition one, participants were asked to attend to the tastiness of the food when making their choices, in the second condition, to attend to the healthiness of the food, and in the third condition, to choose naturally. The instructions emphasized that participants should always choose what they preferred to eat regardless of the attention/consideration cues. Before each block, the attention condition cue was displayed for 5 seconds. On each choice trial, participants had 3 seconds to answer and were shown feedback on their choice for 0.5 seconds after responding. Trials were separated by a variable fixation period of 4 to 6 seconds. Most participants responded on a 4-point scale “strong yes”, “yes”, “no” or “strong no” to indicate if they preferred to eat or to not eat the food shown on the current trial. Five out of 33 participants completed a version of the task including a fifth option that allowed them to signal indifference between eating and not eating the food.

We followed the original analysis procedures in IAC and analysed all 33 subjects as one set. After the scan, participants rated the 180 food items for taste (regardless of health) and health (regardless of taste), with the order of rating types randomized across participants. After both the choice task and ratings were complete, one trial from the choice task was randomly chosen to be realised. Participants were required to eat the food if they answered “yes” or “strong yes”. If they answered “no” or “strong no”, they still had to stay in the laboratory for the 30-minute waiting period; however, they were not allowed to eat any other food. Participants were fully informed of these choice incentivization procedures before beginning the study.

### Data set 4 – Transcranial direct current stimulation study (TDCS)

All participants in this study were pre-screened during recruitment to ensure that they were actively following a healthy lifestyle. They were specifically asked if they would agree to do their best to choose the healthier option whenever possible on the day of the study. Participants who indicated that they would not do so were still allowed to complete the experiment and were reimbursed for their time, but we did not analyse their data. All participants received a flat fee of 100 CHF (approx. 104 USD).

The procedures for this study were originally described in ^72^. We repeat that description here to make this paper self-contained.

#### Participants

The Ethics Committee of the Canton of Zurich approved the study protocol and all participants provided written informed consent. In total, 199 participants were enrolled in the study. No participants reported any history of psychiatric or neurological conditions or had any acute somatic illness. Participants were pre-screened in telephone interviews to ensure they did not suffer from any allergies, food intolerances, or eating disorders. To ensure that the snacks in the food choice task would present a temptation, participants were only eligible if they reported regularly consuming snack foods (at a minimum 2-3 times per week) while at the same time trying to maintain an overall balanced and healthy diet.

Data from 25 participants were excluded because they failed to meet *a priori* inclusion criteria or data quality checks. Within the study we requested a written statement of compliance with a health goal for the time of the experiment (see below). Seven men and 1 woman indicated they would not comply with the health goal; their data were excluded from all analyses. Note that these participants still completed the experimental procedures and received the same compensation through food and monetary incentives as those who complied, so there was no incentive for the participants to lie about following the health goal. Data from 8 participants had to be excluded because they confused the response keys or forgot the identity of the reference item during the task. Four participants were excluded on site due to safety precautions regarding tDCS. Three participants were excluded on site because a re-check of the inclusion criteria revealed that they did not actually like snacks or only consumed them on 1-2 occasions per month instead of the minimum 2 times per week. One additional participant had to be excluded because the choice set could not be constructed due to the fact that he reported only the most extreme values on all health and taste ratings. Lastly, data from one participant was excluded because she never made a healthy choice when taste and healthiness were in conflict in the baseline condition, precluding inference about within-subject changes due to stimulation. This left 87 men and 87 women in the final dataset.

Participants were randomly allocated to stimulation conditions. The anodal (58 participants, 30 female), cathodal (57 participants, 30 female), and sham (59 participants, 27 female) stimulation groups did not differ from each other with regard to age, body mass index (BMI), or self-reported eating patterns (as assessed by the Three Factor Eating Questionnaire, German validated version by Pudel and Westenhöfer ^67^) (see Supplementary Table 9). The groups also did not differ with regard to impulse control (in the stop signal reaction time, SSRT), working memory capacity (digit span test), or time discounting preferences. Finally, the groups did not differ in the level of hunger that they reported before the choice task (see Supplementary Tables 10-17).

#### tDCS stimulation protocol

The target electrode (5 x 7 cm) was placed on the left dlPFC (see Supplementary Figure 2). The reference electrode (10 x 10 cm) was placed over the vertex, off-centred to the contralateral side in such a way that a 5 x 7 cm area of the reference electrode was centred over the vertex while the remaining area was placed more to the right side. The target electrode covered the two dlPFC regions depicted in Supplementary Figure 2 (MNI peak coordinates = [−46 18 24] and [−30 42 24]). These targets were selected because they both showed greater activity for health challenge success > failure in two previous fMRI studies ^52, 54^. The coordinates for both dlPFC and vertex were identified in each participant’s individual T1-weighted anatomical MR image using a neuronavigation system (Brainsight, Rogue Research, RRID:SCR_009539, https://www.rogue-researcher.com/; see Supplementary Figure 2). We applied anodal, cathodal, or sham tDCS over this dlPFC site using a commercially available multi-channel stimulator (neuroConn GmbH). Between a ramp-up and ramp-down phase of 20 seconds, active stimulation with 1 milliampere (mA) took place for 30 minutes (anodal and cathodal group) or 5 seconds (sham). Sham stimulation was delivered with either the anode or the cathode over the dlPFC, counterbalanced over the whole sham group. Both the participants and the experimenters mounting the tDCS electrodes were blind to the stimulation condition.

#### Procedure

Participants first rated 180 food items for health and taste. They were instructed to rate taste regardless of the healthiness and vice versa for each of our 180 food items on a continuous scale that showed visual anchor points from −5 (“not at all”) to +5 (“very much”). Before or after these ratings, participants completed a battery of control tasks in randomized order. All control tasks were performed both before and after stimulation: a stop signal reaction time task (SSRT), a self-paced digit span working memory (WM) test, and a self-paced monetary inter-temporal choice task (ITC). In order to test for stimulation effects on taste and health ratings, participants also re-rated a subset of foods after stimulation (see Supplementary Results).

After all pre-stimulation tasks had been completed, but before any food choices were made, we asked participants to sign a health goal statement in which they indicated whether they would commit to maintaining a health goal during the following food choice task or not (see SI section 1.2 for an English translation of the health goal text). Participants indicated that they would or would not commit to the goal, dated, and signed the document, and then handed it back to the experimenter. Participants could not see which option others in the room had selected and the experimenter randomizing the tDCS conditions was blind to the participants’ responses to the health goal.

Just prior to beginning the food choice task, participants indicated their current hunger levels. They then completed a series of food choices. The first 101 participants made 60 food choices at baseline, however we increased the number of baseline choices to 80 for the final 98 participants in order to have an even number at baseline and under stimulation. All other experimental factors were kept the same for all 199 participants. The baseline choices allowed us to make within-subject comparisons of health challenge success before and during stimulation. Once participants had finished making the baseline choices, stimulation was applied. Participants did not make any choices for the first 3 minutes of stimulation to allow the current to stabilize. Following the stabilization period, they completed another set of food choices (n = 120 for participants 1:101 and n = 80 for participants 102:199). No choice pairs were repeated between the baseline and stimulation choice sets.

However, the difficulty in terms of taste difference was balanced across the two choice sets (see Supplementary Information). Participants completed the set of food choices under stimulation (or sham) in a maximum of 16 minutes. In the remaining 8-14 minutes of stimulation (or sham) time, participants completed several control tasks. We randomized the order of the post-stimulation control tasks so that all tasks had an equal chance of being run in the period when current was still being applied versus the 5-10 minute window immediately after stimulation (during which physiological aftereffects of the tDCS were still present, see ^70, 71^). Once they had completed all post-stimulation control tasks, participants filled in a questionnaire battery (Three Factor Eating Questionnaire (TFEQ), Cognitive Reflection Test (CRT), “Big Five” personality dimensions (NEO-FFI), socio-economic status). They also indicated whether and to what degree they had tried to comply with the health goal throughout the study, whether they had felt the stimulation and how strongly, and whether they had any problems understanding or following the instructions. Finally, participants received and ate their selected food 30 minutes after they made their final decision in the food choice task.

#### Food choice paradigm

Participants were asked to eat a small meal of approx. 400 kcal 3 hours prior to the study and consume nothing but water in the meantime. In the health challenge paradigm, participants chose which food they wanted to eat at the end of the study. In order to comply with their health goal, they had to choose the healthier item as often as they could.

However, the paradigm was engineered such that health and taste of the food options always conflicted based on the participant’s ratings, so they would always have to forgo the tastier food in order to choose healthy. Participants knew that one of their choices would be realised in the end, and they would have to eat whatever they chose on the trial that was randomly selected.

Participants were shown the picture of a reference food for 3 seconds at the beginning of each block. This reference food was either healthier and less tasty than all 10 items shown in the upcoming block or tastier and less healthy than all 10 upcoming items. On each of the 10 trials within a block, participants had to decide if they preferred to eat the food currently shown on the screen or the reference food at the end of the study. The identity of the reference food was written in text on the screen so that participants did not need to remember it (see Figure 1d). During each choice trial, participants had 3 seconds to make their decisions, and each trial was separated by a jittered inter-trial interval of 2-6 seconds. One trial was selected at random to be realised after all experimental procedures were completed. At the end of the study, participants stayed in the lab for 30 minutes to eat the food they obtained in the study.

### Statistical Analyses

For all Bayesian modelling analyses, we used the default, uninformative priors specified by the brms, BEST, or BayesFactor R-packages (see Supplementary Methods). These analyses are not predicated on assumptions of normally distributed data or equal variances across groups. Throughout the paper, the notation PP() indicates the posterior probability that the relation stated within the parentheses is true, similarly the Bayes factors (BF) represent the relative evidence for this relationship over its opposite (e.g. greater than zero versus less than zero). Whenever we analysed previously published data, we applied the same subject- and trial-level exclusion criteria described in the original papers.

### Time-varying Drift Diffusion Model with separate attribute consideration onset times

We fit a drift diffusion model that allowed for differential onset times for taste and health attributes during evidence accumulation to participants choice outcome and reaction time data. Several of the food choice tasks used a 4-point decision strength scale, and for these data we collapsed choices into a binary yes/no or left/right choice. The following six free parameters were estimated separately for each participant and condition:

Thr: evidence threshold for responding (symmetric around zero)

Bias: starting point bias for the evidence accumulation process nDT: non-decision time

RST: relative start time for health (positive values mean that health enters the process after taste, negative values mean health enters before taste)

ω_taste_: weighting factor determining how much taste contributes to the evidence accumulation rate.

ω_health_: weighting factor determining how much healthiness contributes to the evidence accumulation rate.

The values of these six parameters were used to simulate choices and response times using the sequential sampling model described in the equation below to update the relative evidence level at each subsequent time step *t*.

If taste enters first, the update equation is

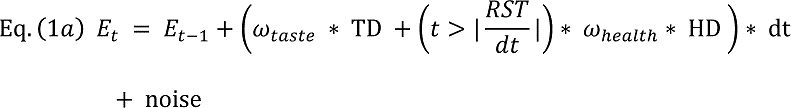

While if healthiness enters first it is

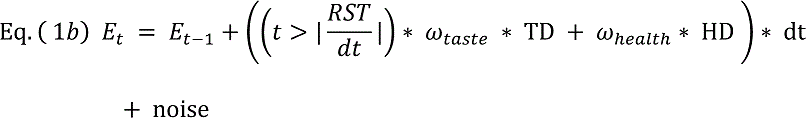

Thus, the times at which the weighted value differences in tastiness and healthiness attributes (ω_taste_*TD and ω_health_*HD, respectively) begin to influence the evidence accumulation rate are determined by RST. When the conditional statement 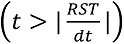 is false, it equals 0, while if true it equals 1. Multiplying one of the two weighted attribute values by 0 until 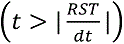 is true means that that attribute does not factor into the evidence accumulation process for the initial time period determined by |RST|. The RST parameter is defined as the consideration start time for healthiness minus the starting time for tastiness. Thus, RST will have a positive value when tastiness enters into consideration first and a negative value when healthiness is considered first. Note that the standard, synchronous onset DDM is equivalent to the specific case of *RST* = 0, and then equations (1a) and (1b) are equivalent because *t* is always greater than (|*RST* / dt|) and 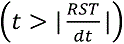 always equals 1.

Based on previous mouse-tracking results from Sullivan et al. ^16^, our model formulation makes the simplifying assumption that once an attribute comes into consideration it continues to influence the rate of evidence accumulation until the choice is made. Model comparisons testing this assumption showed that the different starting time formulation fit the data better than DDMs that allowed for differential attribute consideration end times, both different start and end times, non-decision times that varied as a function of choice outcome, or starting point biases in favour of tastiness (see Figures 2, 3, Extended Data Figure 2, and Supplementary Table 1).

Evidence accumulation proceeds according to equation (1a) or (1b) in the following manner. The evidence accumulation process begins with an initial value (E_0_) that is equal to the value of the Bias parameter. This value is then updated in discrete time-steps of dt = 0.008 s until |E_t_| is greater than the threshold (Thr) parameter value. The noise at each step of the accumulation process is drawn from a Gaussian distribution with mean = 0. The differences in taste and healthiness ratings between Food1 and Food2 (or Food1 vs. 0 for the single item choices in the IAC dataset) on a given trial are denoted by TD and HD, respectively. Once the threshold is crossed, the response time is computed as t*dt + nDT, where nDT is a free parameter for a non-decision time that accounts for the time required for any initial perceptual or subsequent motor processes that surround the period of active evidence accumulation and comparison.

We estimated the best values for all six free parameters described above separately for each participant and condition using the Differential Evolution algorithm described in Mullen, et al. ^73^ with a population size of 60 members run over 150 iterations. On every iteration, we simulated 3000 decisions and response times for all unique combinations of taste and healthiness trade-offs in the participant’s choice set using each population member’s six tDDM parameters. We then computed the likelihood of the observed data given the distribution generated by the 3000 simulated choices for a given set of parameters. On each subsequent iteration, the population evolves toward a set of parameters that maximize the likelihood of the observed data using the procedures described by Mullen and colleagues ^73^.

We examined the evolution of the population over the 150 iterations/generations and found that the Differential Evolution algorithm settled on a set of best-fitting parameters well before 150 iterations in our data sets. The upper and lower bounds on the search space for each of the 6 parameters are listed in Supplementary Table 18. The ratings for taste and healthiness were z-scored across all available ratings of each type for the whole set of participants in each study.

Lastly, we also fit a standard DDM and several other alternative DDM formulations (see Supplementary Table 1) to all datasets using the same procedures as the tDDM. The simulated choice sets from each model shown in Figure 2 were composed of 1000 repetitions of each of the 673 unique taste and health combinations used across all four data sets.

We also note that we fit the tDDM using two levels of resolution for the tastiness and healthiness ratings in the GFC and TDCS studies. The tastiness and healthiness ratings from these two studies were collected on a 426-point visual analogue scale. We initially fit the tDDM using the 426-point ratings scale. We also estimated the fits after first reducing the resolution to 10 equally-sized bins (i.e., 42.6 points per bin) for both taste and health. Both versions yielded very similar results, but the estimation proceeded considerably faster when using the binned ratings because this reduced the number of unique combinations of attributes and therefore the number of simulations required for the fitting procedure. We report the parameter values and results from the model with binned ratings for these studies.

### Tests of parameter recovery

We generated simulated choices and reaction times by parameterizing the standard and tDDMs using the best fitting parameters for each model estimated from the choices made by participants in the baseline condition for all four studies. The simulated choice sets were based on these parameters and the tastiness and healthiness differences participants faced on every decision trial. Thus, the simulated choice sets matched the empirical data in terms of trial numbers and attribute difference distributions. Fitting these simulated choices allowed us to quantify both models’ ability to recover known parameter values within the context of our experimental datasets and the ability to distinguish between these models (see Extended Data Figure 1, Supplementary Figure 1 and Supplementary Table 1).

### Testing taste versus healthiness influence by response time

In addition to parameter recovery tests, we used simulated choices to test how well each model reproduced choice and response-time characteristics observed in the empirical data. A hierarchical Bayesian logistic regression analysis showed that the influence of taste and healthiness on choice outcomes differed as a function of response times. (equation (2); Tables S2-4). More specifically, this analysis tested the influence of each attribute on trials in which the response was made before versus after the relative-starting-time advantage of the first attribute had elapsed, on average. The population-level regressors are listed in equation (2) below.

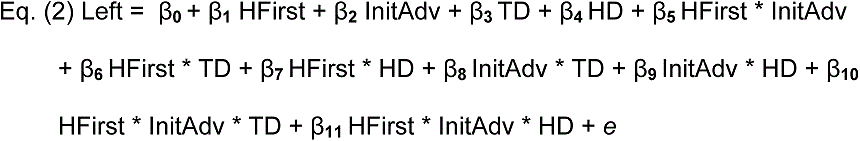

In this equation, Left is a binary indicator of the choice outcome. HFirst is a dummy variable (1 = healthiness, 0 = taste) indicating which attribute is considered first – as determined by the tDDM. InitAdv is a dummy variable (1 = before, 0 = after) indicating whether the response was made before the median value of the sum of relative-starting-time difference plus non-decision time across participants had elapsed. This sum was equal to 1 second. One second was also the cut-off for first quartile of the response time range, meaning that 25% of choices were made in 1 second or less. The abbreviations TD and HD stand for the differences in tastiness and healthiness, respectively, on each trial. Subject-specific coefficients were estimated for all regressors except HFirst, because each participant had only one level of that regressor in his or her in baseline condition.

We computed this regression using four different subsets of empirical or simulated data. Initially, we analysed the baseline choice trials pooled over all 4 studies (baseline = mouse response trials from MRT, day 1 trials from GFC, no-cue trials from IAC, pre-stimulation trials from TDCS). We then compared this model to two simpler models that omitted either 1) the dependency on response time (i.e. InitAdv dummy variable) or 2) both the dependency on response time and the indicator for which attribute a participant considered first (i.e. InitAdv and HFirst dummy variables). The full model explained the data better (see Supplementary Table 4), and therefore, we used it to examine choice patterns generated by the standard and tDDM. The means and 95% HDIs for regression coefficients plotted in Figure 3a come from estimating the hierarchical logistic regression in equation (2) to observed, time-varying (tDDM), standard (sDDM), or tastier starting-point-bias (bDDM) model-simulated choices for all participants in whose |RST| parameter fell into the third quartile. We subset the data into this quartile so that timing differences between taste and healthiness would be big enough to have a clear effect in both the real and simulated data.

### Correspondence of tDDM health delay estimates with MRT estimates

With their mouse response trajectory analysis, Sullivan and colleagues were able to estimate to within a fraction (1/101) of each response time when health first became and remained significant in each choice (their Figure 4b). In order to compare our estimate (which was given in seconds and represents a mean value across all of a given set of choices) to the MRT estimates, we transformed the MRT estimates of start times for health into a mean estimate in seconds as well. Specifically, we took the mean of the estimated trial-wise health start time bins for each participant and multiplied it by the participant’s mean RT, then divided by 101. The MRT method was only able to estimate health start times for N=18 (out of 28) participants and, therefore, we calculated the Bayesian equivalent of Pearson’s correlation coefficient between tDDM and mouse-tracking estimates of health start times in this subset of participants. Unless otherwise noted, all correlation coefficients reported in this paper represent the mean of the posterior distribution from a Bayesian correlation analysis. These Bayesian correlations were implemented in R and JAGS based on code published on the blog, doingbayesiandataanalysis.blogspot.com, that accompanies the “Doing Bayesian Data Analysis” book by Kruschke ^74^.

### Relationship between relative-start-times and other tDDM parameters

To explain how individual differences in the relative-start-time for healthiness were related to the other tDDM parameters (Supplementary Table 6), we estimated the model specified in equation (3) below:

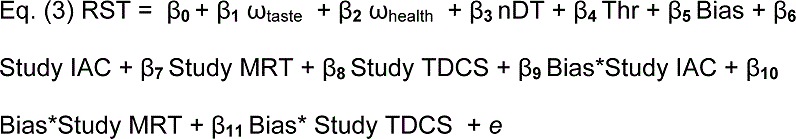

Note that we interacted the Bias parameter from the tDDM with a dummy variable indicating the Study, because the bias measures different answers across studies given the task designs (e.g. left/right, eat/do not eat). The GFC study served as the baseline in this regression.

### Out of sample tests for comparing the standard and tDDMs

We fit the standard and tDDM with separate attribute consideration onsets to the odd-numbered choices from each participant and then compared the accuracy of the two models when predicting even-numbered choice outcomes. We used the squared error in predicting choice outcomes as our measure of accuracy ^75^. The predicted outcome for each choice was computed as the mean outcome over 1000 simulations from the standard and tDDM. Choices for the food on the left or to eat the food in single option decisions were set to a value of 1, and the alternative choice was set to a value of 0. Thus, the mean outcome from the 1000 simulations for each choice represented the probability of a given outcome. The scoring rule for accuracy on each trial was then computed as: (True_Outcome – Prediction)^2^. We computed the squared error separately for tastier and less tasty choice outcomes and then took the mean error across these trials types to obtain a measure of balanced error.

### Changes in tDDM parameters between instructed attention conditions

We compared tDDM parameters fit to choices during health-cued (HC), taste-cued (TC), and natural-cued (NC) blocks using a Bayesian t-like test (implemented in the R Package, BEST version 3.1.0 ^34^), which in turn relies on JAGS (version 3.3.0).

### Modelling changes in behaviour under tDCS

We first fit the hierarchical regression model specified in equation (4) to the odd-numbered baseline trials in our tDCS dataset. Based on those fitted parameters, we generated predictions about the probability of health challenge success in even-numbered trials as a function of tDCS polarity (anodal, cathodal, sham), stimulation session (baseline, active), health difference, taste difference, and participant identity. We then estimated equation (4) on all even-numbered trials for which the probability of health challenge success was predicted to be between 0.2 and 0.8.

To examine whether stimulation over left dlPFC caused changes in health challenge success, we fit a Bayesian hierarchical logistic regression model to the tDCS dataset. The population-level regressors for this model are given in condensed notation in equation (4).

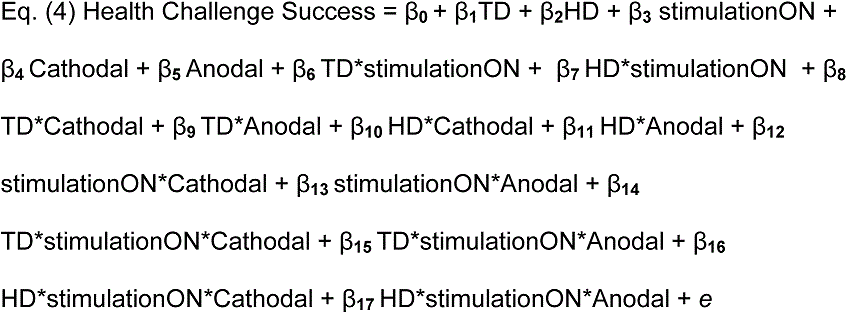

Here, TD and HD denote the absolute value of taste and healthiness difference between foods on each trial, stimulationON was a dummy variable taking the value 1 under stimulation and 0 at baseline, and Anodal and Cathodal were the active stimulation conditions. The Sham condition was the baseline in this regression. The model included the main effects of all regressors as well as the two and three-way interactions between attribute differences and stimulation Type and session (i.e., baseline vs. stimulation on). The model also included subject-specific intercepts, stimulation effects, and slopes for HD and TD (Supplementary Table 8).

### Choice of sample sizes, randomization, and blinding

Studies IAC and MRT are based on pre-existing datasets and we included the participants analysed in the original papers. No statistical methods were used to pre-determine sample sizes for the GFC study and TDCS studies but our sample sizes are similar to those reported in relevant previous publications^18, 52, 76^. In the TDCS study, participants were assigned randomly to the stimulation condition. Experimental conditions in the TDCS and GFC studies were blocked. In the TDCS study, the baseline condition was always presented first in order to prevent any influence of potential tDCS after-effects. The food choice and gambling conditions in GFC study were presented in random order. Data in the TDCS study was collected double-blind. Data analyses were performed with knowledge of the condition labels in all experiments.

### Data availability

The data analysed in this paper are openly available on the Open Science Framework at, https://osf.io/g76fn/. Additional data for the mouse response trajectory experiments from Sullivan et al. (2015) are available at https://osf.io/jmiwn/

### Code availability

The code for fitting the diffusion models and running the other analyses is openly available on the Open Science Framework at, https://osf.io/g76fn/.

## Acknowledgements

This study has been supported by Swiss National Science Foundation grants 320030_143443 and 32003B_166566 (CCR and TAH), a Marie Curie International Incoming Fellowship PIIF-GA-2012-327196 (ARB), and EU FP7 grant 607310 (TAH). The funders had no role in the conceptualization, design, data collection, analysis, decision to publish, or preparation of the manuscript. The authors thank Aidan Makwana and Ana Cubillo for contributing code for the food choice and control tasks, Jonathan Price for help with data collection and documentation, and Gaia Lombardi for help with data collection and documentation as well as useful discussions about analysis strategies and implementation.

## Author contributions

S.U.M., A.R.B., R.P., C.C.R. and T.A.H. designed one or more aspects of the research; S.U.M. and A.R.B. collected the novel data in studies GFC and TDCS; R.P. and T.A.H. designed the time-varying drift diffusion model with separate attribute consideration onset times; S.U.M., A.R.B., R.P., and T.A.H. analysed the data; S.U.M., A.R.B., R.P., C.C.R. and T.A.H. wrote the paper.

## Supplementary Information

### Supplementary Results

#### Model comparison and parameter recovery tests for the standard DDM and time-varying DDM with separate attribute consideration onset times

The time-varying DDM with separate consideration onset times for taste and healthiness (tDDM) yielded a better fit to choice and reaction time distributions across all subjects (N = 272) than the standard formulation of a DDM with a single, synchronous onset time (Extended Data Figure 1; tDDM BIC = 280632, standard DDM BIC = 281909). Furthermore, the improvement in the fit for the tDDM over the standard DDM is proportional to the absolute value of the estimated relative start time for a given participant (Supplementary Table 19). This relationship is expected because the greater the difference in onset times between taste and healthiness, the more behaviour will deviate from the predictions of the standard, synchronous onset DDM.

We also tested the out-of-sample accuracy for standard versus tDDM fits to our human participants’ choices. Specifically, we fit the standard and tDDMs to the odd-numbered choices from each participant and then quantified the accuracy of each model when predicting even-numbered choice outcomes. The mean squared error for the tDDM (0.163) was lower than that of the standard DDM (0.170; posterior probability tDDM error < sDDM error = 0.97). Supplementary Table 2 shows the results of a hierarchical Bayesian linear regression on the trial-wise improvement in prediction error for the tDDM over the standard DDM as a function of the taste and healthiness differences in each choice and the best fitting tDDM parameters for each human participant. Thus, overall the tDDM better explains response-time dependent influences of taste and healthiness on choice outcomes, response time distributions regardless of whether the choice was healthy or unhealthy, and makes more accurate out-of-sample predictions for choice outcomes than the standard DDM and several other DDM specifications suggested by anonymous reviewers (see Supplementary Table 1).

We performed parameter recovery tests for both the standard and tDDM. These tests were based on the actual food choices each of the 272 participants faced in the lab experiments. Specifically, we simulated one set of choices using the best-fitting standard DDM parameters and a second set using the best-fitting tDDM parameters for each participant using the choice options (i.e. taste and healthiness differences) the participant faced in the lab. We then fit both simulated choice sets using the standard and tDDM. This gave us four new sets of fitted parameters that we could compare with the known generating parameters and models.

Parameter recovery was very good for both the standard and tDDM when fitting to the corresponding generating model (Extended Data Figure 1 and Supplementary Figure 1; Supplementary Table 20). Furthermore, when fitting the tDDM to choices generated by the standard DDM, there was no correlation between errors in the relative weights versus errors in the relative timing (mean r = 0.02, PP (r < 0) = 0.36, 95% HDI = [−0.09; 0.15], Bayes factor (BF) = 0.192; Extended Data Figure 1). However, when fitting the standard DDM to choices generated by the tDDM, the errors in relative weight versus relative timing parameter recovery were negatively correlated (mean r = −0.44, PP (r < 0) > 0.99995, 95% HDI = [−0.54; −0.35], Bayes factor > 1000). This pattern of biased errors is expected because a standard DDM is forced to account for the difference in attribute consideration onset timing by over-estimating the difference in weights between the two attributes.

In addition to these parameter recovery tests, we performed model identification/recovery tests on simulated data. At the model level, we were able to determine whether the generating model was the standard or a time-varying DDM with separate consideration onsets for taste and healthiness in each case. These recovery tests were based on out-of-sample predictive accuracies. To compute the out-of-sample prediction accuracy we generated 1000 choices for 150 new food pairs from 272 real-subject-based simulated agents that used either the generating or recovered standard and tDDM parameters. We then quantified out-of-sample accuracy by computing the squared error between new choices (i.e. the mean left/right outcome over 1000 simulated choices for each food pair) made by the original generating parameters and recovered standard or tDDM parameters. When the tDDM was the generating model, the recovered tDDM parameters had lower squared error (mean = 0.006) than the recovered standard DDM parameters (mean = 0.016). In contrast, when the standard DDM was the generating model, the recovered tDDM parameters had higher squared error (mean = 0.012) than the recovered standard DDM parameters (mean = 0.008). Supplementary Table 21 shows the results of a hierarchical Bayesian regression on the trial-wise improvement in prediction error for the tDDM over the standard DDM as a function of the taste and healthiness differences in each choice and the simulated agents’ tDDM parameters. Note that, in this regression on simulated data, the included tDDM parameters are the ones known to have generated the choices.

### Results from control analyses in the tDCS study

#### TDCS effects in other cognitive domains

We assessed potential differences in several cognitive domains under tDCS using the Bayesian regression model specified in Supplementary Equation (S1) in the supplementary methods. We found no effects of tDCS stimulation over left dlPFC on working memory (Supplementary Tables 10-11), response inhibition (Supplementary Tables 12-13), or monetary inter-temporal choice (Supplementary Tables 14-15). The paradigms used to measure these behaviours are described in the supplementary methods below. Note that although previous papers have reported effects of brain stimulation over the dlPFC in several of these domains, our stimulation target and electrode placement is different from those previous studies.

#### Pre-choice hunger levels

To assess the hunger level, we report percentages of maximum hunger level as participants indicated it on a visual analogue scale (Supplementary Table 16). Self-reported hunger did not differ between the stimulation groups (for results of the Bayesian regression model specified in Supplementary Equation (S2) see Supplementary Table 17).

### Stability of taste and health ratings before and after stimulation

In order to test the stability of taste and health ratings after stimulation, participants re-rated a random subset of 30 foods drawn from the original set of 180 foods after stimulation. We tested the stability of both rating types with Bayesian linear models specified as in Supplementary Equation (S3) in the Supplementary Methods below. For the health ratings, neither stimulation session (mean beta estimate = 1.11 ± 0.75 SD, 95% credible interval: [−0.34; 2.58]), nor stimulation condition (estimate = −0.43 ± 0.82, CI: [−2.05;1.20]), nor their interaction (estimate = −0.13 ± 0.34, CI: [−0.82; 0.54]) showed significant differences at the population level (Supplementary Table 22a). Taste ratings were slightly higher for all groups after stimulation (estimate = 3.70 ± 0.95, CI: [1.86; 5.57]), and higher at baseline for the anodal (estimate = 4.92 ± 1.92, CI: [1.14; 8.71]) and cathodal groups (estimate = 4.18 ± 1.92; CI: [0.42; 7.91]) compared to sham, but the critical interaction test showed that stimulation groups did not differ in their taste rating increase between pre- and post-stimulation measurements (estimate of the stimulation session X condition interaction = −0.16 ± 1.31, CI: [−2.75; 2.41] for cathodal and for anodal estimate = −1.07 ±1.31, CI: [−3.69; 1.49]; Supplementary Table 22b). Note that the ratings were scaled to fall between 0 and 100, and thus the coefficients can be interpreted as percentage of the visual analogue rating scale.

### No relationship between RST and SSRT and WM in the baseline trials from the tDCS study.

There was no correlation between the start time for health relative to taste (RST) estimated during baseline (i.e., without stimulation) choice trials and inhibitory control, measured as stop signal reaction time (mean r = 0.05, PP (r >0) = 0.71, 95% HDI = [−0.12; 0.233], BF= 0.342), or RST and working memory capacity, measured as forward digit span (mean r = 0.02, PP (r > 0) = 0.61, 95% HDI = [−0.125; 0.17], BF = 0.221).

### Testing the relationship between attribute rating response times and relative-start-times during choice

We reasoned that rating response times might be an index of how readily taste and healthiness attributes come to mind and play a role in determining their relative onset times. Therefore, we tested whether the time required to report taste and healthiness ratings during the separate rating sessions was associated with the relative-start-times for these attributes during the decision process (see Supplementary Equation (S4) in the supplementary methods). Using the data from the baseline session in our tDCS experiment (i.e., the largest sample in one homogenous experimental setup), we found that the time it took to report taste or healthiness ratings was not significantly associated with relative-start-times (taste regression coef. = 0.05 ± 11 sec, 95% HDI = [−0.17; 0.28], health regression coef = 0.07 ± 0.12 sec, 95% HDI = [−0.17; 0.31]; Supplementary Table 23).

## The influence of attribute consideration onset times on choice outcomes and response times

We also ran two hierarchical Bayesian regressions showing how relative-start-times (RSTs) relate to both response times and choices. The time-varying DDM with separate attribute consideration onset times predicts that the influence of RST on a given choice will depend on the individual’s subjective weights for tastiness and healthiness. Furthermore, it also predicts that the influence of RST on choice outcomes will depend on whether or not the tastiness and healthiness attributes both favor the same option or they are in conflict (i.e. differs between health challenge and non-challenge trials). The regressions in Supplementary Tables 24 and 25 show that these predicted interactions are present in our data.

## Supplementary Discussion

We used the DDM as a starting point for our modelling analysis because this flavour of sequential sampling model is relatively simple, well established, and widely used to fit choice and response-time data across cognitive domains.

However, a number of different sequential sampling model formulations exist, and in specific cases, these models make different predictions about choice and reaction-time distributions ^1–7^. However, in our food choice datasets, most of these sequential sampling models will be nearly indistinguishable ^8–10^; we therefore refer to our current model as a sequential sampling model to emphasize that we are adding a feature to one representative of this larger class of models. Adding this flexibility for attributes to enter into consideration at different points is also possible for many other sequential sampling models. We also note that our results from the tDDM are consistent with theoretical and empirical work showing that sequential sampling models, including drift diffusion models and linear ballistic accumulators, can capture changes in perceptual decision processes that result from known changes in externally presented evidence over time ^6, 11–14^. However, in contrast to previous work on perception, we tested for asynchronous attribute consideration onsets in value-based choices for which the externally presented evidence is constant. In other words, we examined timing differences resulting from internal cognitive and neural processes instead of changes in the stimuli themselves. Regardless of the context (e.g. perceptual, value-based) or the exact form of sequential sampling, the key feature that allows these models to explain the data is the ability to account for variability in the strength of evidence over time.

We also note that an attribute-specific, independent race model is not a parsimonious explanation of our data. In an independent race model, the first competitor to cross the threshold determines how the decision is made ^15^.

Therefore, in a race model that included separate and independent accumulators for each attribute value or difference in attribute values, the response times will only be proportional to the value of the winning attribute (i.e. the one that crosses the threshold first). This prediction of race models does not hold in our data. Regardless of whether or not participants chose the healthier or tastier option, response times are significantly related to the difference in both attributes (Supplementary Tables 26 and 27). Moreover, response times also depend on whether or not the subjective value derived from the initially considered attribute alone is compatible (i.e. points to the same choice) with a subjective value computed by integrating across both taste and healthiness attributes (see Supplementary Table 28 and Figure 3). Thus, our results are more consistent with hierarchical comparisons of attributes on one level and integrated option values at a second level, as in Hunt and colleagues’ fMRI study of risky choices ^1^. There must be integration across attributes at some level in the evidence accumulation hierarchy or mutual inhibition between attribute-specific accumulators to explain the response time results we find.

The use of analysis strategies that quantify the separate effects on relative timing and weighting, such as the tDDM, is important for interpreting brain stimulation data on the role of dlPFC and other brain regions in value-based choices. Previous studies have reported that stimulation targeted over various brain regions causes changes in several forms of decision making including choices over trade-offs between monetary amounts and risk or time, or between rewards for oneself and others ^16–24^. Notably, all of these choices involve multi-attribute stimuli and, frequently, conflict between the different attributes. In light of our modelling results, we can speculate that the mechanistic change caused by stimulation over the dlPFC is in the attribute weighting process in some cases. However, the different studies have targeted a range of dlPFC coordinates across both the left and right hemispheres and have used various forms of brain stimulation with potentially different local and widespread effects. Therefore, one should not assume that altered attribute weighting is the mechanistic result of every dlPFC-targeted stimulation protocol. Fortunately, asynchronous evidence accumulation modelling methods, such as the separate attribute consideration onset tDDM used here, could be applied to most of the existing datasets cited above or newly acquired data to gain further insights into how and why brain stimulation causes changes in choice behaviour. Moreover, such analyses are by no means limited to brain stimulation and can be applied to any set of response-time and choice data on multi-attribute decisions (e.g. self/other, amount/delay, risk/magnitude) under different biological or experimental conditions. This may elucidate other neural regions that are involved in determining the relative timing of attribute consideration.

## Supplementary Methods

### TDCS control task descriptions

The following procedures were originally described in ^27^. We repeat that description here to make this paper self-contained.

## Stop signal reaction time task (SSRT)

We used a standard stop-signal-reaction time task ^28–31^ in order to assess whether stimulation changed inhibitory control. Participants had to press a button as quickly as they could whenever a figure appeared on the screen (“go task”), but had to stop the initiated movement if another figure appeared above the first with a few milliseconds delay (“stop signal”). The initial delay between the stop signal and the go signal was 0.25 seconds. The task was adaptive and added 0.05 seconds delay to the next inhibition trial whenever the participant’s rate of successful movement inhibition was greater than 50% over all previous inhibition trials (adding up to a delay of maximum 0.95 seconds), and subtracted 0.05 seconds whenever the participant’s success rate in inhibiting the button press fell below 50% over all inhibition trials completed up to this point. Stimuli were presented on the screen with a jittered duration between 0.5 and 1.25 seconds, and late responses that were given after the stimulus had disappeared from the screen were not counted as correct. Trials with (25) and without stop signal (75) were randomly mixed within the run.

The stop signal reaction times (calculated by subtracting the average presented delays from the average reaction times) were negative for a number of participants, indicating that they adopted a strategy of waiting for the stop signal to appear before initiating a response. In other words, they did not follow the task instructions to initiate a response as soon as the go signal appeared on the screen. We removed participants showing this pattern from all analyses using SSRTs. The remaining sample included 39 Anodal, 41 Cathodal and 42 Sham participants.

## Digit span task

In order to test and, if necessary, account for changes in working memory capacity under tDCS, participants completed a computerized digit span task according to the procedure of Wechsler ^32^. The screen first showed a series of 5 numbers, each for 1 second, and then prompted the participant to enter the numbers as she remembered them. If the participant entered a correct sequence in ascending (“forward”) order two times in a row, the difficulty level increased by one digit (up to a maximum of 12 digits). If the participant failed two times in a row, or alternated between correct and incorrect answers more than seven times without reaching two sequential correct responses, the task stopped and prompted participants to enter the digits in reverse of the order they were originally shown (“backward”) in the next round of trials. Within the backwards digit span trials, participants also needed two consecutive correct responses to reach the next level, stopped at 12 digits, or if they reached the failure criteria described above. We excluded data from two participants for the following reasons: For one participant in the Sham group, data for the post-stimulation control were lost due to a computer crash. One participant in the Anodal group was detected to cheat by writing down all sequences on the instruction sheet. We report data from 172 participants (57 Anodal, 57 Cathodal and 58 Sham). Note that due to an oversight in the task programming, it was possible to cheat and solve the backward digit span by entering the numbers in the forward direction (i.e., the program did not force participants to enter responses in the requested order). We therefore only evaluated the forward digit span.

## Inter-temporal choice task

To test and, if necessary, account for possible effects of tDCS on discounting behaviour, we ran an inter-temporal choice (ITC) task based on the paradigm of Cooper, Kable, Kim & Zauberman ^33^. Participants were instructed that they would earn part of their total payment in this task (60 CHF were paid as a baseline on the day of the study, and the present discounted value of 40 CHF from the ITC was paid at the time specified by the participant). They were told that we would randomly draw select one trial from either the baseline or stimulation session and realize (i.e., pay out) their decision on that trial. In this version of the ITC task, participants participated in a BDM auction ^34^ designed to elicit their indifference points between a payoff on the day of the experiment and 40 CHF at various points in the future. Participants were asked to specify a bid between 1 and 40 CHF for which they would be indifferent between receiving the bid amount today versus 40 CHF after the specified delay on that trial. Participants made bids for immediate payoffs versus 40 CHF in 14 linearly spaced delays ranging from 13 to 181 days from the day of the experiment.

The rules of the BDM auction were fully explained to participants and were as follows. If the participant’s bid was more than a randomly determined counter offer that was uniformly drawn from the range 0:40 CHF, she would receive the delayed payment of 40 CHF in the indicated number of days. However, if she had bid less than the counter offer, she was paid the amount of the counter offer at the day of the study instead of 40 CHF in the future. If the bid was equal to the counter offer, then a coin flip decided which payment was made. This mechanism ensures that it is in the participants’ best interest to bid their true value for the present equivalent of 40 CHF at the given delay.

We calculated a discounting score for each participant as the area under the curve for all bids, where higher bids indicate a greater willingness to wait for the delayed outcome. We excluded the data from several participants from analysis because their bidding patterns indicated that they did not understand the task (e.g., they alternated between bidding high and low amounts with increasing delay, or bid lower amounts for short delays and increased their bids for longer delays, which is the opposite of the expected pattern). The ITC data analyses we report are based on N = 135 participants (45 Anodal, 45 Cathodal, and 45 from the Sham group) who showed a consistent pattern of temporal discounting across trials.

## Regression models used in the tDCS study

All analyses were conducted using Bayesian Markov-chain Monte Carlo (MCMC) methods with R ^35^, in combination with STAN ^36^ or JAGS ^37^. We used the default, uninformative priors specified by the brms ^38^ or BEST ^39^ R-packages.

We tested working memory, monetary intertemporal choice and SSRT task performance (Supplementary Tables 11, 13, and 15) as a function of stimulation session and stimulation condition using linear regressions taking the form of equation (S1):

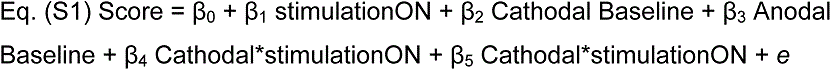

where Score denoted the respective working memory, intertemporal choice or stop-signal reaction time performance, stimulationON was a binary dummy variable denoting stimulation session (0 = pre-stimulation, 1 = concurrent or immediately after stimulation), and Anodal and Cathodal were the active stimulation conditions. The Sham stimulation condition served as the baseline in this regression.

Baseline differences in hunger levels (Supplementary Table 17) were modelled according to Supplementary Equation (S2):

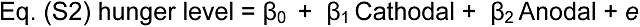

where hunger level was given in percent (as indicated on a visual analogue scale with 0% not at all and 100% = maximally hungry). The Sham stimulation condition served as the baseline in this regression.

The population-level regressors for the taste and health ratings regressions (Supplementary Table 22) are listed in Supplementary Equation (S3) below. Note that the models also included intercepts for subjects and food items (i.e., each subject and food item was treated as a random effect) as well as subject- and food-specific specific slopes for the effect of stimulation.

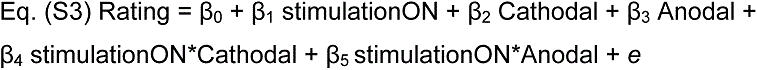

where Rating was either health or taste ratings for the foods (tested in separate models), stimulationON was a factor for stimulation session (pre-stimulation, post-stimulation), and Anodal and Cathodal were the active stimulation conditions. The Sham condition was the baseline in this regression.

## Health goal statement used in the tDCS study

The statement read: “In this study, we want to investigate how people make healthy food choices. Therefore, we ask you to maintain the goal of eating as healthy as possible during this study. Specifically, we ask you to try and choose the healthier of the two food options on each trial. However, these are real decisions, and you are required to eat the food that you chose in one randomly selected trial. We realize this may be more difficult for some people than others, and it is important for us to know whether you agree to this goal or not. Your participation and payment are not contingent on your response. However, this is important for the scientific validity of our study, so please mark your answer below honestly. Please mark “yes” if you agree to do your best to follow the health goal. Please mark “no” if you do not want to commit yourself to the health goal.” Participants made their selections, dated and signed the document, and handed it back to the experimenter.

## Comparing response times during rating sessions to relative start times

We used response times during the rating sessions as an estimate of the participants’ fluency in recalling or constructing taste and healthiness attributes (Supplementary Table 23). To test whether the relative start time (RST) depended on the speed of ratings for either health or taste aspects, we estimated the following model for each participant:

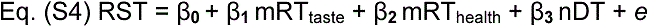

where mRT is the mean reaction time over all taste ratings or health ratings that the participant made at the beginning of the experiment, and nDT is the non-decision time estimated in the tDDM. We conducted this analysis using the data from the baseline session in our tDCS experiment (i.e., our largest set of data from a single choice paradigm/context).

## Testing the influence of all tDDM parameters on the improvement in fit for the tDDM relative to the standard DDM

We tested how the improvement in fit for the tDDM relative to the standard DDM (Supplementary Table 19) relates to all the parameters of the tDDM by estimating the following Bayesian linear model:

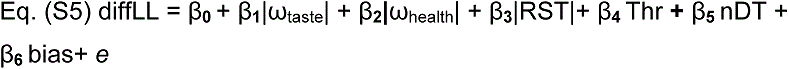

where diffLL is the difference in Log-Likelihood between the tDDM and the standard DDM, |ω_taste/health_| is the absolute value of taste or the health weights, |RST| is the absolute value of relative-start-time (to accommodate changes in any direction), Thr is the threshold, and nDT is the non-decision-time estimated by the tDDM. We conducted this analysis using models fit to the choice and reaction time distributions across all participants (N = 272).

Note that similar results are obtained when using a logistic regression that treats the difference in Log-Likelihood as a binary outcome.

## Improvement of prediction accuracy for the tDDM relative to standard DDM on choices made by a tDDM generating process and by human participants

We tested how much the out-of-sample prediction accuracy increased for a time-varying relative to a standard DDM on choices made by human participants (Supplementary Table 2) as well as on choices made by simulated agents that used a time-varying DDM to make decisions (Supplementary Table 21). We modeled this improvement as a function of trial and individual-specific variables using the following Bayesian hierarchical linear regression:

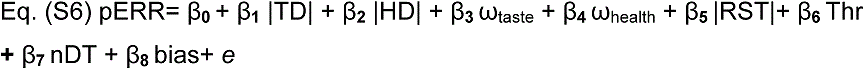

where pERR indicates the percent error reduction of the tDDM over the standard DDM in predicting out-of-sample choices, |TD |and |HD| are the absolute trial-wise health and taste value differences, and ω_taste/health_, Thr, nDT, bias, and |RST| indicate the 6 tDDM parameters: taste and health weighting, threshold, non-decision time, bias, and the absolute value of the relative start time for health.

## Testing for differences in the strength of association between relative attribute weights and starting times in generating versus recovered parameters

We tested whether there was a difference in the strength of association between relative attribute weights and starting times in recovered relative to known generating parameters. The recovered parameters were estimated from choices generated by the best fitting tDDM parameters across 272 participants (Supplementary Table 20). We conducted this analysis using the following Bayesian linear model:

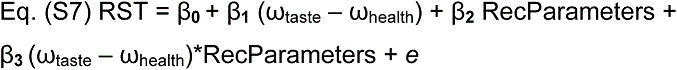

where RST refers to the relative starting times across generating and recovered parameters, (ω_taste_ – ω_health_) are the relative attribute weights, and RecParameters is a dummy variable where 1 or 0 indicates that the parameter was recovered or generated, respectively.

## Testing healthy food choices as a function of attribute weighting and timing parameters

We modeled healthy food choices as a function of the estimated parameters of the tDDM, the difference in taste and healthiness ratings, and whether the participants faced a challenge on the respective trial by fitting a Bayesian hierarchical logistic regression (Supplementary Table 24). The population-level regressors are given below:

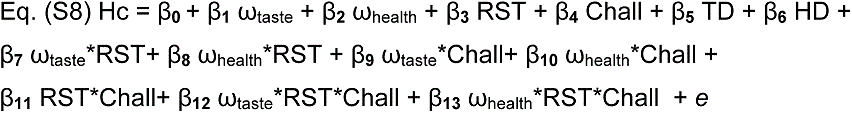

where Hc is a binary variable indicating whether a healthy food item was chosen, ω_taste/health_ are the estimated taste or health weighting parameters of the tDDM, RST is the relative start time for health, Chall is a dummy variable indicating whether the participants faced a challenge on the respective trial, and TD and HD are the differences in taste and healthiness ratings between the two options, respectively. The model included subject-specific intercepts as well as subject-specific slopes for TD and HD to capture individual differences in sensitivity to these two aspects when making healthy choices.

## Testing response times in health challenge and non-challenge trials as a function of the tDDM parameters

We used a hierarchical Bayesian linear regression to model response times in health challenge and non-challenge trials as a function of the tDDM parameters pooled across the baseline conditions of all 4 studies (Supplementary Table 25). The population-level regressors are indicated below:

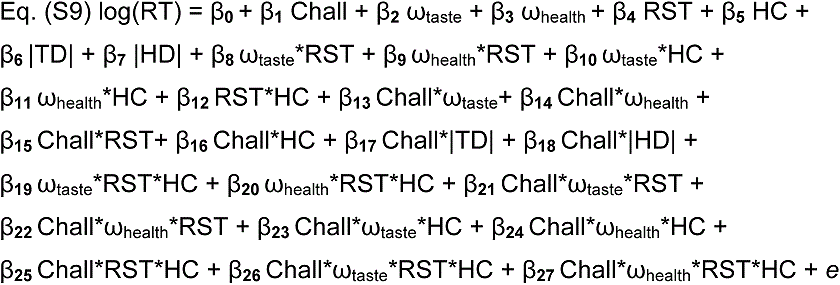

where log(RT) are log-transformed reaction times, Chall is a dummy variable indicating whether the participants faced a challenge on the respective trial, ω_taste/health_ are the estimated taste or health weighting parameters of the tDDM, HC is a binary variable indicating whether a healthy food item was chosen, and |TD| and |HD| are the absolute value differences in taste and healthiness ratings between the two options, respectively. The model included subject-specific intercepts and subject-specific slopes for the |TD| and |HD| regressors as well as the interactions of these regressors with the challenge trial and healthy choice dummy variables to capture individual differences in sensitivity to taste and health aspects when making healthy or unhealthy choices in challenging or non-challenging settings.

## Modeling response times in cases where the healthier or tastier option was chosen

We tested how response times were affected by the difference in attribute ratings and whether participants faced a challenge on the respective trial separately for cases where the healthier (Supplementary Table 26) or tastier option was chosen (Supplementary Table 27) with the following regression:

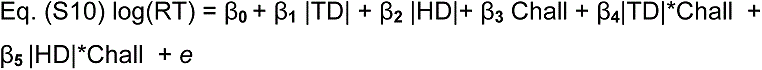

where log(RT) are log-transformed reaction times,|TD| and |HD| are the absolute value differences in taste and healthiness ratings between the two options, respectively, and Chall is a dummy variable indicating whether the participants faced a challenge on the respective trial. The model included subject-specific intercepts and subject-specific slopes for the |TD| and |HD| regressors and their interaction with the challenge trial dummy variable to capture individual differences in sensitivity to taste and health aspects in challenging or non-challenging settings.

## Modelling response times as a function of subjective value when one or two attributes are considered

To test whether response times also depend on whether or not the subjective value derived from the initially considered attribute alone is compatible (i.e. points to the same choice) with a subjective value computed by integrating across both taste and healthiness attributes (Supplementary Table 28), we modeled response times with the following regression:

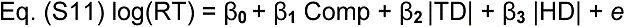

where log(RT) are log-transformed reaction times, Comp is a dummy variable indicating whether the subjective value on the respective trial for the initially considered attribute was compatible with the subjective value of both health and taste attributes combined, and |TD| and |HD| are the absolute value differences in taste and healthiness ratings between the two options, respectively. The model included subject-specific intercepts as well as subject-specific slopes for |TD| and |HD| to capture individual differences in sensitivity to these two attributes.

## Statistics packages used

All analyses presented in this paper were performed with the R ^35^, STAN ^36^ and JAGS ^37^ statistical software packages. The sequential sampling model was fit using the Rcpp toolbox ^40^. Bayesian regression models were run using the brms package ^38^ which is an interface between R and STAN. All correlations and t-like tests were computed using Bayesian MCMC sampling methods using R in combination with JAGS ^39, 41^. Bayes factors for the correlations and t-like tests were computed using the BayesFactor ^42^ package in R. The plots in Extended Data Figure 3 and Figures 4 and 5 were created using the yarrr package ^43^. The packages pracma ^44^ and plyr ^45^ were used for data handling and restructuring.

## Extended Data Figures

**Extended Data Figure 1.**
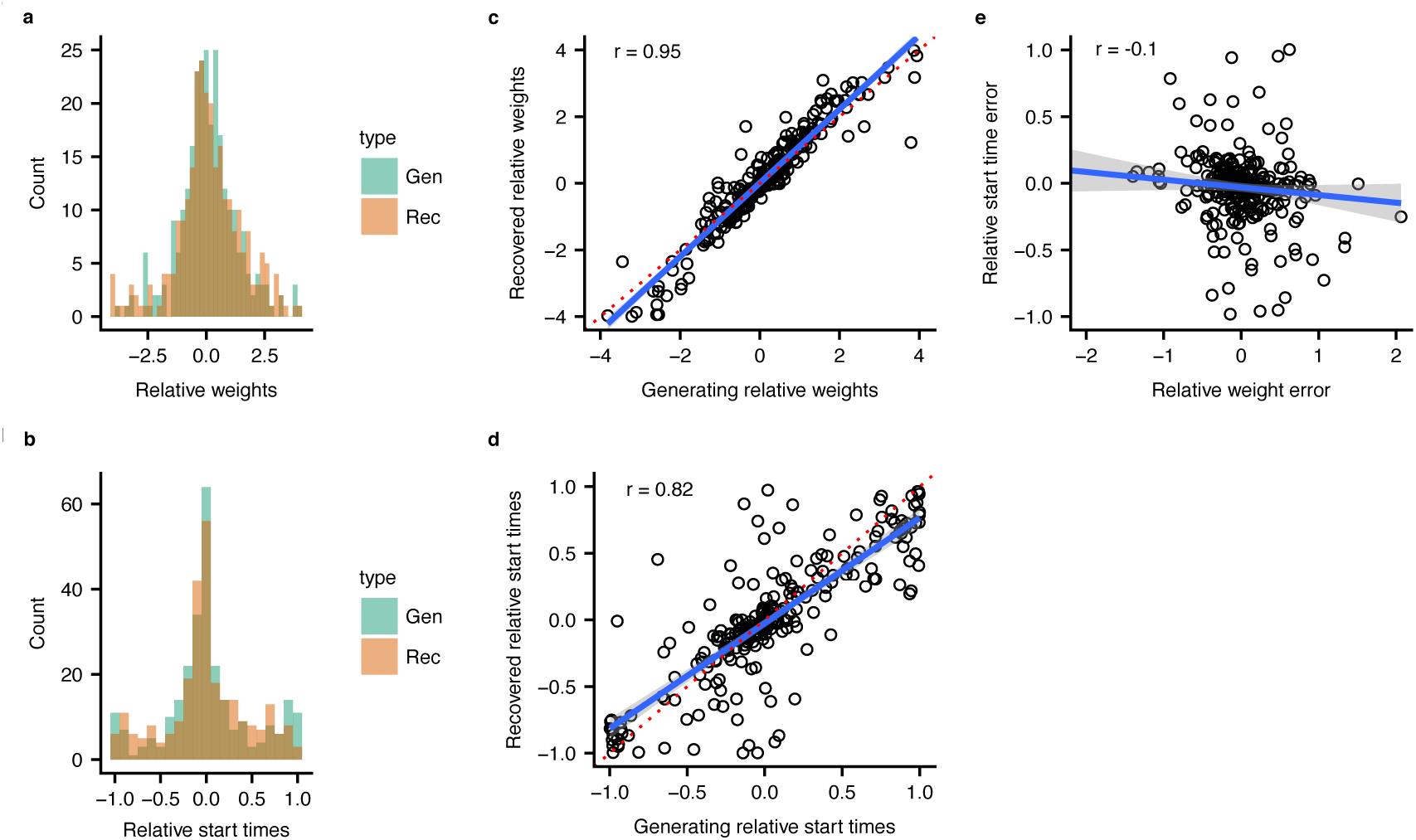
Parameter recovery for the time-varying DDM with separate consideration onset times for tastiness and healthiness attributes. The plots in the first column show the distributions of all 272 generating and recovered relative weighting (a) and timing parameters (b). There was no significant difference between generating and recovered relative weighting (mean difference = 0.01, 95% HDI = [−0.36, 0.54], posterior probability of a difference > 0 = 0.662, Bayes factor = 0.140) or relative timing parameters (mean difference = −0.01, 95% HDI = [−0.03, 0.01], posterior probability of a difference > 0 = 0.105, Bayes factor = 0.024). The panels in the second column show the correlations between the generating and recovered relative weighting (c) and timing parameters (d). The red dotted line indicates the x = y identity line. Panel e) plots the error in relative weight recovery against the error in relative timing recovery. This plot shows that there is no significant correlation between the two types of error when fitting the model (r = −0.1, 95% HDI = [−0.215; 0.018], posterior probability of observing a negative correlation = 0.95). The grey shaded area (panels c-e) signifies the 95% confidence interval.

**Extended Data Figure 2.**
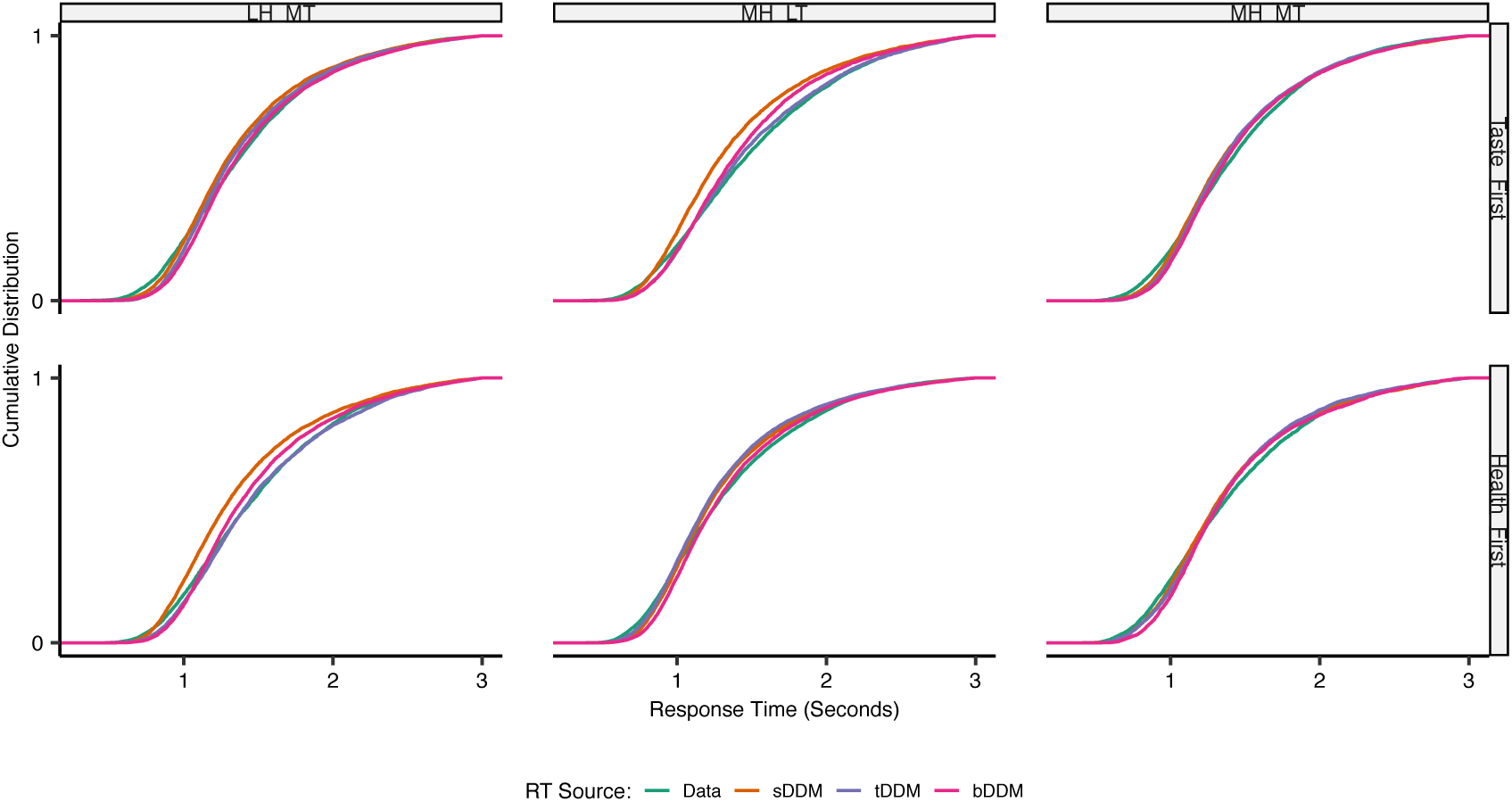
Cumulative distributions for response times by participant type, choice outcome and data source. Participants estimated to consider taste or health first are plotted in the top and bottom rows, respectively. Response times for choices in favour of 1) less healthy but more tasty (LH_MT), 2) more healthy but less tasty (MH_LT), or 3) both more healthy and more tasty (MH_MT) outcomes are shown in columns 1-3, respectively. Choices in favour of the option rated as less healthy and less tasty were rarely made (less than 5% of trials) and are omitted for clarity. Responses generated by human participants are shown in green lines. Responses generated by simulated agents using the best-fitting sDDM, tDDM, and bDDM parameters are shown in orange, purple, and magenta lines respectively. All three models can recreate the RT patterns in the empirical data equally well when choice outcomes align with the attribute participants consider first. However, the sDDM and bDDM both generate response times that are too fast relative to the empirical data when participants that consider taste first ultimately choose in favour of a more healthy, but less tasty option (row 1, column 2) or if participants that consider health first ultimately choose in favour of a less healthy, but more tasty option (row 2, column 1). In contrast, the tDDM is able to reproduce the observed response time distributions in these cases well.

**Extended Data Figure 3.**
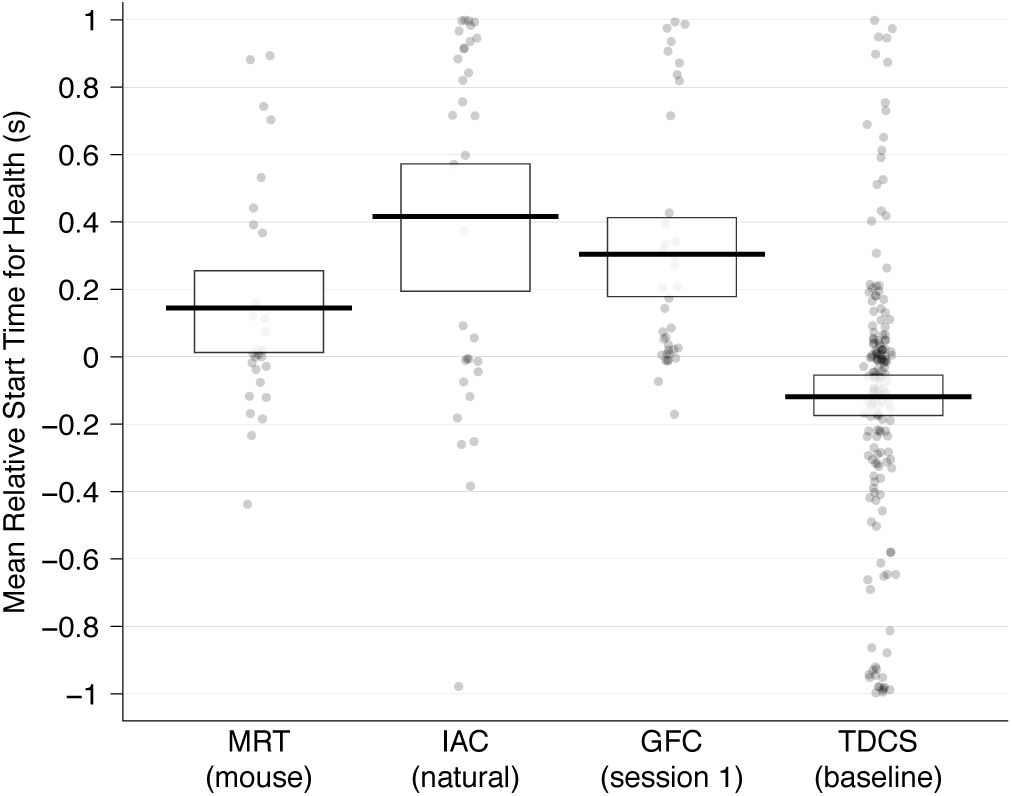
Relative start times in seconds for healthiness compared to tastiness for all participants in each study. Positive values indicate that tastiness is considered before healthiness and negative values that healthiness is considered before tastiness. In each column every dot is a separate participant. The thick black horizontal bars represent within-study means and the rectangular bands indicate the 95% highest density intervals (HDIs). Dataset abbreviations: MRT = data from the computer-mouse response trials in Sullivan et al 2015; IAC = data from the natural choice condition in Hare et al 2011; GFC = newly collected data from the first session/day of an experiment combining gambles and food choices; TDCS = newly collected data from the pre-stimulation baseline choices in our tDCS experiment.

**Supplementary Figure 1.**
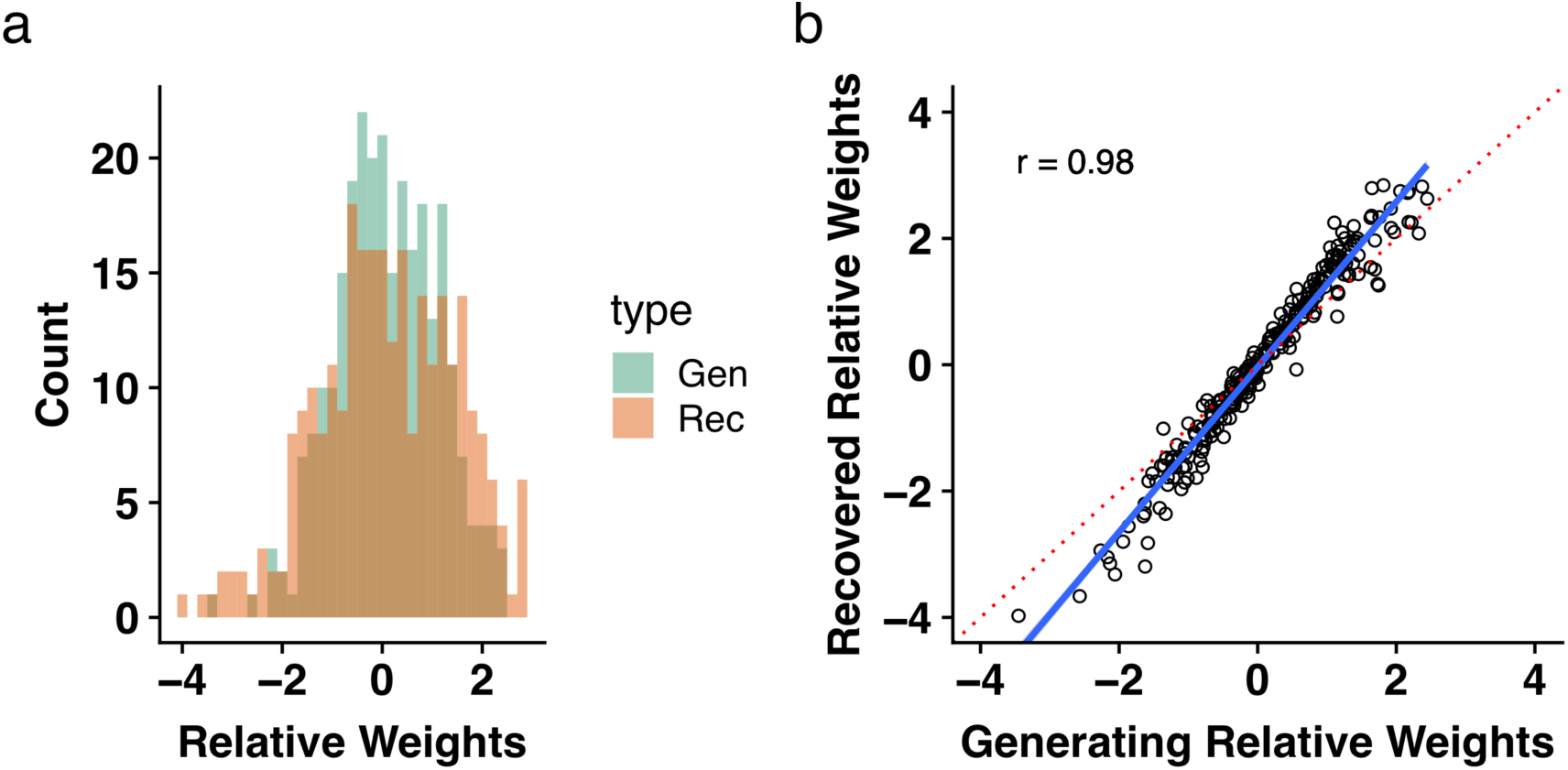
Parameter recovery for the standard DDM with simultaneous consideration onset times for tastiness and healthiness attributes. **a)** shows the distributions of all 272 generating and recovered relative weighting parameters. Recall that the relative consideration start time is fixed at zero for the standard DDM. There was no significant difference between generating and recovered relative weighting (mean difference = 0.004, 95% HDI = [−0.05, 0.05], posterior probability of a difference > 0 = 0.58, BF = 0.061). **b)** shows the correlation between the same generating and recovered parameters from the histogram in panel **a**. The red dotted line indicates the x = y identity line.

**Supplementary Figure 2.**
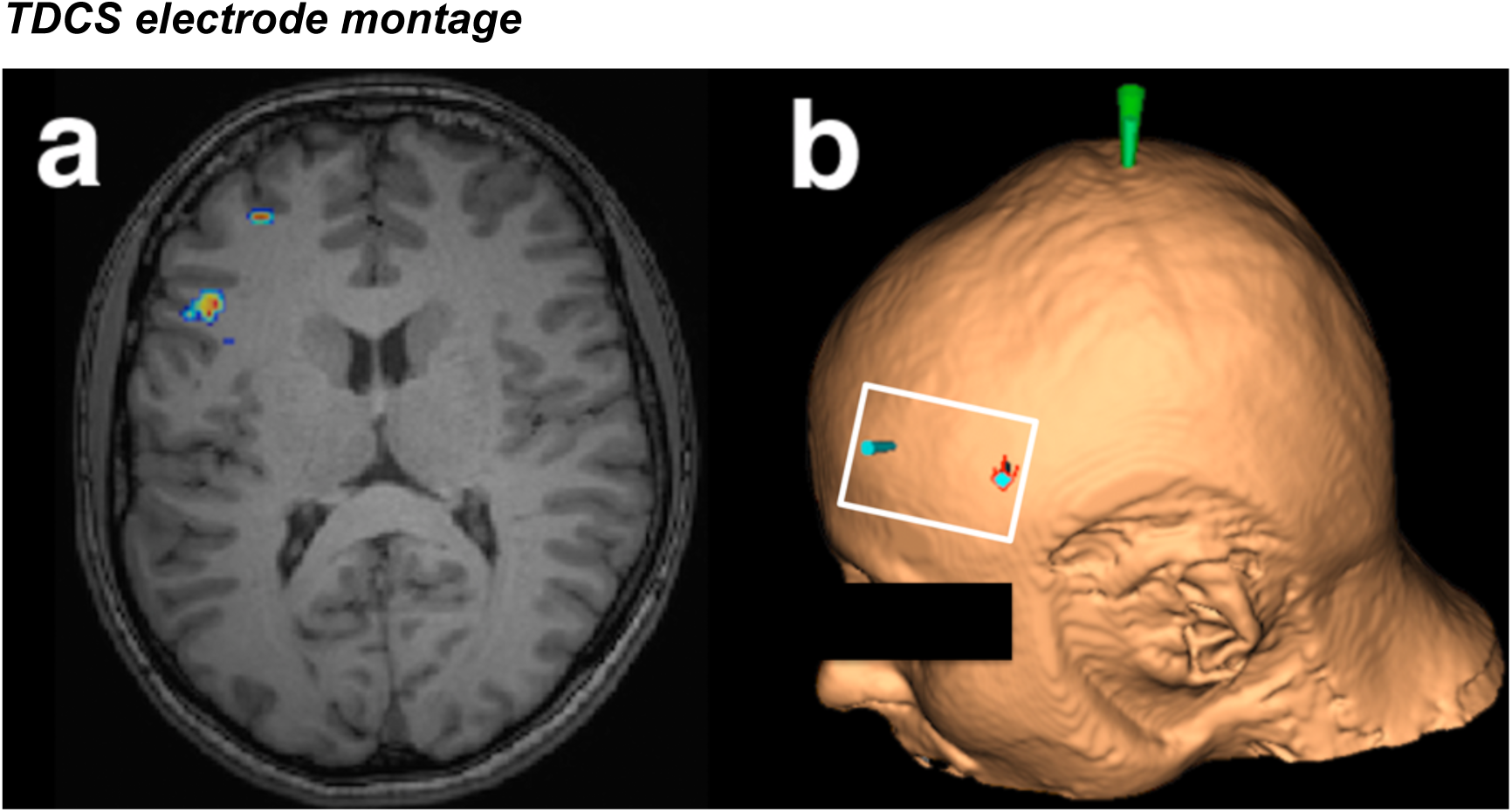
Left dlPFC target regions in the tDCS study. **(a)** The stimulation was centred over two previously identified dlPFC regions of interest (MNI peak coordinates = [−46 18 24] and [−30 42 24]) based on contrasts of health challenge success > failure in two previous fMRI studies (Hare, et al. ^25^ and Maier, et al. ^26^). **(b)** The dlPFC and vertex coordinates were identified with neuronavigation based on anatomical brain scans for each participant. A 5×7 cm electrode (indicated by the white frame) was placed over left dlPFC to cover both stimulation targets, and a 10 x 10 cm reference electrode was placed over the vertex slightly offset to the right hemisphere (so that the centre of a comparable 5×7 cm area would be centred over the meeting point of the two central sulci, see Methods).

### Supplementary Tables

**Supplementary Table 1.** Model fit metrics for alternative specifications of the DDM.

| DDM type | Squared Error |  | BIC |  |
| --- | --- | --- | --- | --- |
|  | Squared Error | Difference | BIC | difference |
| **Different Onsets** | 0.163 | - | 280632 | - |
| **Standard DDM** | 0.17 | 0.007 | 281909 | 1277 |
| **Different Offsets** | 0.168 | 0.005 | 286493 | 5861 |
| **Different Onsets & Offsets** | 0.187 | 0.024 | 318724 | 38092 |
| **Starting-point bias for tastier** | 0.168 | 0.005 | 282311 | 1679 |
| **Variable non-decision time** | 0.304 | 0.141 | 290650 | 10018 |

This table lists the squared error in out-of-sample predictions and the Bayesian information criterion (BIC) values for the time-varying DDM used throughout the main text, the standard DDM, and three other alternative DDM formulations. Column 1 lists the squared error in predictions for even-numbered trial outcomes generated from parameter fits to participants' odd-numbered choices. Column 2 lists the increase in error for a given model relative to the tDDM with separate consideration onset times for taste and healthiness attributes. Thus, higher values indicate better performance for our primary tDDM. Column 3 lists the BIC values for each model across all trials from all 272 participants in the four data sets we examine here. Column 4 lists the differences between each alternative and our primary tDDM specification (i.e. alternative BIC – primary BIC). Here again, higher values indicate better performance for our primary tDDM. Note that the BIC metric factors in how well the models capture both decision outcomes and response times in-sample, while the squared error metric is based on out-of-sample decision outcomes alone.
Model names and descriptions:
**Different Onsets** = a time-varying DDM that includes a free parameter allowing for the consideration onset time of each attribute to differ. This is the model we use throughout the rest of the paper.
**Different Offsets** = a DDM that instead of different onsets allows for the time at which each attribute ceases to be considered to differ. In other words, one attribute could stop adding to the evidence accumulation process before a threshold is reached.
**Different Onsets & Offsets** = a DDM including both separate onset and offset times for each attribute.
**Starting-point-bias for tastier** = a standard DDM with simultaneous attribute on and offset times. However, in this case the data have been transformed such that tastier outcomes are always on the upper boundary of the DDM. This allows the starting point bias that formally measured left/right or eat/don't eat biases in our other formulations of the DDM to capture a bias in favour of the tastier option.
**Standard** = a standard DDM with boundaries and starting-point biases for left/right or eat/don't outcomes.
**Variable non-decision time** = a standard DDM with simultaneous attribute on and offset times. However, in this case the non-decision time is allowed to vary between 1) trials in which healthiness and tastiness attributes are aligned (i.e. there is no health challenge), 2) health challenge trials in which the participant made the healthier choice, and 3) health challenge trials in which the participant made the tastier choice.
In every case, we fit the DDMs to either all (for BIC calculations) or the odd-numbered choices (for out-of-sample squared error) in one experimental condition per participant. In other words, there are no repeated within-subjects measures in these comparisons. All DDMs were fit using the same procedures as those described in the main-text Methods section.

**Supplementary Table 2.**
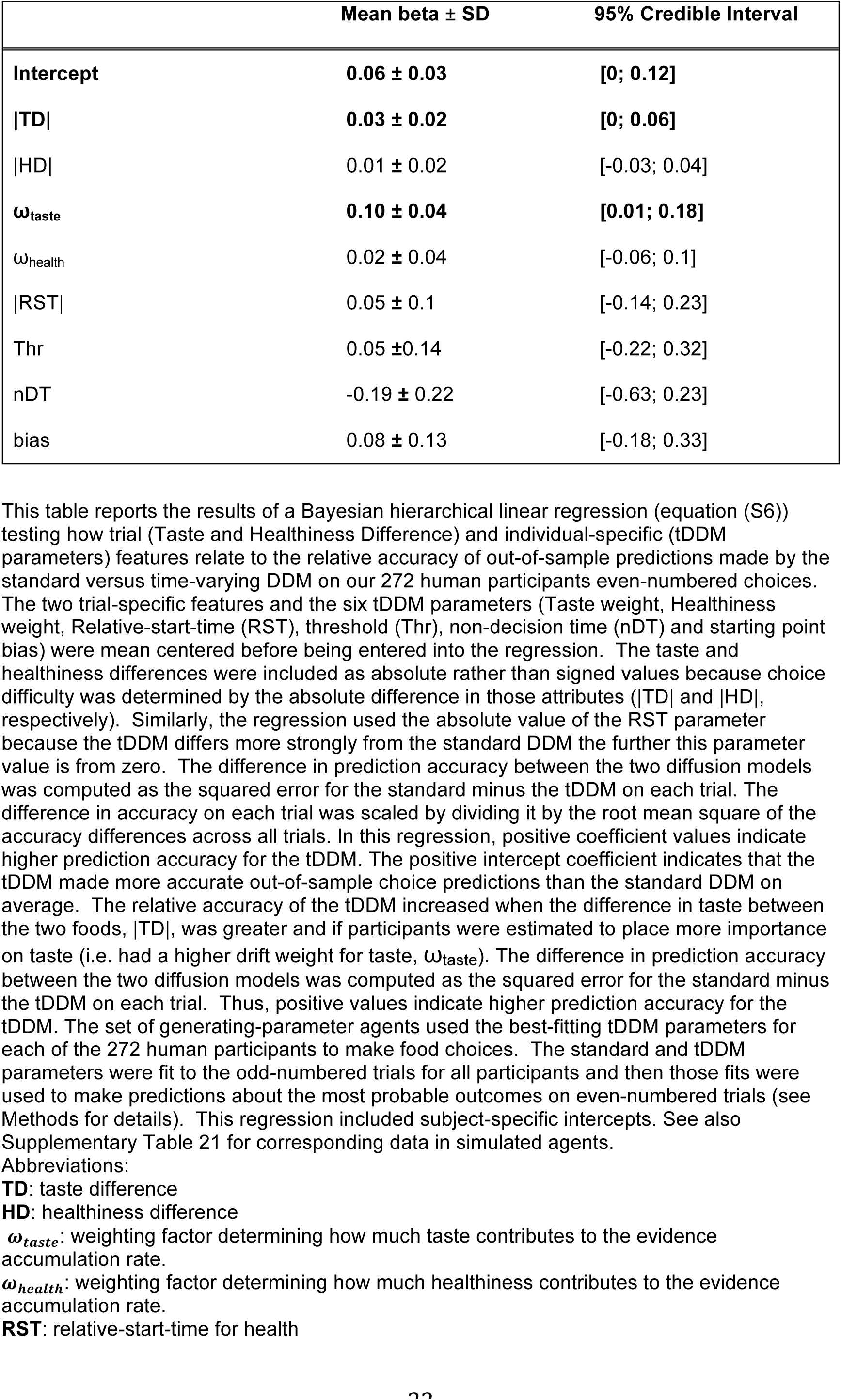
Improvement in out-of-sample prediction accuracy for the time-varying relative to the standard DDM on choices made by human participants.

**Supplementary Table 3.**
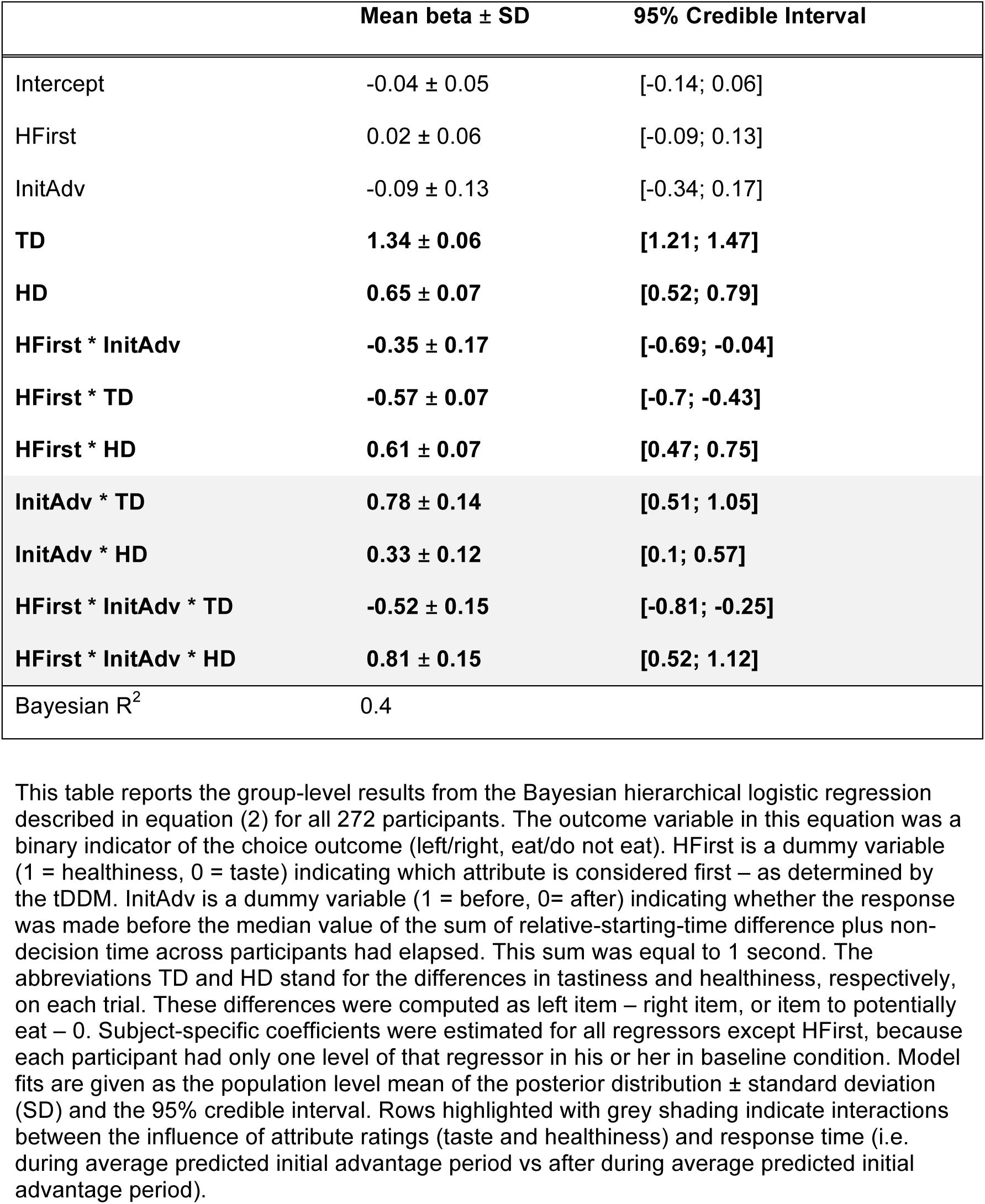
The influence of tastiness relative to healthiness before and after the average initial advantage for the attribute considered first has elapsed.

| | Mean beta ± SD | 95% Credible Interval |
| --- | --- | --- |
| Intercept | -0.04 ± 0.05 | [-0.14; 0.06] |
| HFirst | 0.02 ± 0.06 | [-0.09; 0.13] |
| InitAdv | -0.09 ± 0.13 | [-0.34; 0.17] |
| **TD** | **1.34 ± 0.06** | **[1.21; 1.47]** |
| **HD** | **0.65 ± 0.07** | **[0.52; 0.79]** |
| HFirst * InitAdv | -0.35 ± 0.17 | [-0.69; -0.04] |
| HFirst * TD | -0.57 ± 0.07 | [-0.7; -0.43] |
| HFirst * HD | 0.61 ± 0.07 | [0.47; 0.75] |
| InitAdv * TD | 0.78 ± 0.14 | [0.51; 1.05] |
| InitAdv * HD | 0.33 ± 0.12 | [0.1; 0.57] |
| HFirst * InitAdv * TD | -0.52 ± 0.15 | [-0.81; -0.25] |
| HFirst * InitAdv * HD | 0.81 ± 0.15 | [0.52; 1.12] |
| Bayesian R<sup>2</sup> | 0.4 |  |

This table reports the group-level results from the Bayesian hierarchical logistic regression described in equation (2) for all 272 participants. The outcome variable in this equation was a binary indicator of the choice outcome (left/right, eat/do not eat). HFirst is a dummy variable (1 = healthiness, 0 = taste) indicating which attribute is considered first – as determined by the tDDM. InitAdv is a dummy variable (1 = before, 0 = after) indicating whether the response was made before the median value of the sum of relative-starting-time difference plus non-decision time across participants had elapsed. This sum was equal to 1 second. The abbreviations TD and HD stand for the differences in tastiness and healthiness, respectively, on each trial. These differences were computed as left item – right item, or item to potentially eat – 0. Subject-specific coefficients were estimated for all regressors except HFirst, because each participant had only one level of that regressor in his or her in baseline condition. Model fits are given as the population level mean of the posterior distribution ± standard deviation (SD) and the 95% credible interval. Rows highlighted with grey shading indicate interactions between the influence of attribute ratings (taste and healthiness) and response time (i.e. during average predicted initial advantage period vs after during average predicted initial advantage period).

**Supplementary Table 4.** Model comparisons between a Bayesian hierarchical logistic regression including response time and consideration order effects and two simplified models without those features.

| Model | LOOIC | SE |
| --- | --- | --- |
| Full Model | 22873.33 | 161.16 |
| Reduced Model 1 | 23168.74 | 161.25 |
| Reduced Model 2 | 23342.44 | 159.83 |
| Full – Reduced 1 | -295.41 | 33.46 |
| Full – Reduced 2 | -469.12 | 43.48 |

This table shows the expected out-of-sample prediction error (LOOIC) for three Bayesian hierarchical logistic regressions that measure the relative influence of health and tastiness attributes on choice outcomes. The LOOIC values were computed using Pareto smoothed importance-sampling leave-one-out cross-validation<sup>46,47</sup>. Smaller values indicate better expected out-of-sample performance. The standard errors (SE) for the LOOIC or differences in LOOIC are shown in the last column.
The full model is given by equation (2) that we repeat here for convenience:

**Supplementary Table 5.** The influence of taste relative to healthiness before and after the average initial advantage for the attribute considered first has elapsed in simulated choices.

|  | Empirical data |  | tDDM simulations |  | sDDM simulations |  | bDDM simulations |  |
| --- | --- | --- | --- | --- | --- | --- | --- | --- |
| | Mean ± SD | 95% CI | Mean ± SD | 95% CI | Mean ± SD | 95% CI | Mean ± SD | 95% CI |
| Intercept | 0.12 ± 0.12 | [-0.12; 0.36] | 0.07 ± 0.10 | [-0.11; 0.27] | 0.07 ± 0.07 | [-0.07; 0.21] | -0.01 ± 0.12 | [-0.25; 0.21] |
| HFirst | -0.29 ± 0.16 | [-0.61; 0.02] | -0.22 ± 0.13 | [-0.47; 0.04] | -0.24 ± 0.1 | [-0.43; -0.05] | -0.2 ± 0.17 | [-0.53; 0.13] |
| InitAdv | **0.87 ± 0.31** | **[0.29; 1.51]** | **1.67 ± 0.77** | **[0.21; 3.24]** | **0.83 ± 0.43** | **[0; 1.71]** | 0.5 ± 0.43 | [-0.32; 1.35] |
| TD | **1.33 ± 0.14** | **[1.06; 1.60]** | **1.75 ± 0.19** | **[1.38; 2.12]** | **1.23 ± 0.14** | **[0.95; 1.51]** | **1.1 ± 0.15** | **[0.81; 1.39]** |
| HD | 0.29 ± 0.17 | [-0.05; 0.63] | 0.45 ± 0.25 | [-0.03; 0.93] | 0.23 ± 0.16 | [-0.08; 0.53] | 0.22 ± 0.14 | [-0.05; 0.5] |
| HFirst * InitAdv | **-1.73 ± 0.42** | **[-2.59; -0.95]** | **-3.62 ± 1.01** | **[-5.81; -1.81]** | **-1.43 ± 0.54** | **[-2.54; -0.41]** | **-2.16 ± 0.62** | **[-3.47; -1.04]** |
| HFirst * TD | **-0.50 ± 0.18** | **[-0.86; -0.15]** | **-0.77 ± 0.24** | **[-1.24; -0.29]** | **-0.58 ± 0.19** | **[-0.96; -0.2]** | **-0.44 ± 0.19** | **[-0.81; -0.06]** |
| HFirst * HD | **0.94 ± 0.23** | **[0.50; 1.38]** | **1.28 ± 0.33** | **[0.65; 1.95]** | **1.05 ± 0.21** | **[0.65; 1.48]** | **0.71 ± 0.19** | **[0.34; 1.08]** |
| InitAdv * TD | **1.09 ± 0.30** | **[0.56; 1.73]** | 0.37 ± 0.36 | [-0.29; 1.13] | 0.04 ± 0.2 | [-0.32; 0.45] | 0.37 ± 0.32 | [-0.23; 1.02] |
| InitAdv * HD | -0.06 ± 0.24 | [-0.53; 0.42] | **-0.62 ± 0.31** | **[-1.29; -0.08]** | -0.18 ± 0.17 | [-0.51; 0.17] | -0.01 ± 0.21 | [-0.43; 0.41] |
| HFirst * InitAdv * TD | **-1.31 ± 0.33** | **[-2.01; -0.70]** | **-1.39 ± 0.48** | **[-2.38; -0.51]** | 0.04 ± 0.25 | [-0.46; 0.51] | 0.25 ± 0.42 | [-0.59; 1.04] |
| HFirst * InitAdv * HD | **1.04 ± 0.39** | **[0.37; 1.85]** | **1.31 ± 0.50** | **[0.43; 2.41]** | 0.2 ± 0.25 | [-0.27; 0.68] | 0.53 ± 0.31 | [-0.08; 1.19] |

**Supplementary Table 6.** The variability in relative-start-times is only modestly related to other tDDM parameters.

| | Mean beta ± SD | 95% Credible Interval |
| --- | --- | --- |
| Intercept | 0.19 ± 0.24 | [-0.29; 0.65] |
| $\omega_{\text{taste}}$ | **0.15 ± 0.04** | **[0.07; 0.22]** |
| $\omega_{\text{health}}$ | 0.01 ± 0.04 | [-0.07; 0.08] |
| nDT | 0 ± 0.19 | [-0.37; 0.38] |
| Thr | -0.04 ± 0.11 | [-0.26; 0.18] |
| Bias | -0.27 ± 0.82 | [-1.90; 1.29] |
| Study IAC | 0.07 ± 0.10 | [-0.12; 0.27] |
| Study MRT | -0.10 ± 0.11 | [-0.32; 0.11] |
| **Study TDCS** | **-0.32 ± 0.09** | **[-0.49; -0.15]** |
| Bias X Study IAC | 0.39 ± 0.85 | [-1.22; 2.07] |
| Bias X Study MRT | -0.18 ± 0.92 | [-1.92; 1.65] |
| Bias X Study TDCS | 0.74 ± 0.83 | [-0.86; 2.38] |
| Bayesian $R^2$ | 0.3 | |

**Supplementary Table 7.**
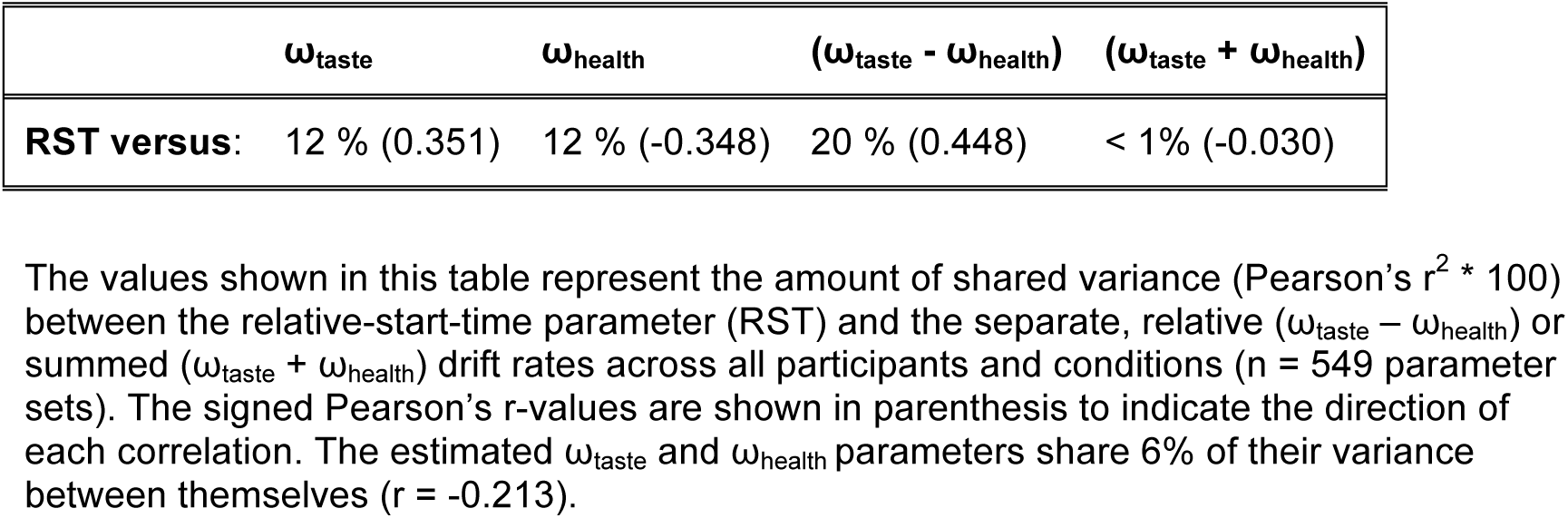
Shared variance between relative start-times and drift weights.

**Supplementary Table 8.** Logistic regression testing whether stimulation over left dlPFC causes changes in health challenge success.

| Model parameter | Mean beta ± SD | 95% Credible Interval |
| --- | --- | --- |
| *Main effect on health challenge success* |  |  |
| Intercept | -0.04 ± 0.12 | [-0.28; 0.21] |
| **TD** | **-0.61 ± 0.11** | **[-0.83; -0.40]** |
| **HD** | **0.72 ± 0.11** | **[0.51; 0.94]** |
| stimulationON | 0.03 ± 0.11 | [-0.18; 0.24] |
| Cathodal | -0.10 ± 0.17 | [-0.44; 0.24] |
| Anodal | -0.02 ± 0.18 | [-0.37; 0.32] |
| TD X stimulationON | -0.02 ± 0.11 | [-0.23; 0.20] |
| **HD X stimulationON** | **0.24 ± 0.11** | **[0.04; 0.45]** |
| TD X Cathodal | 0.08 ± 0.16 | [-0.22; 0.39] |
| TD X Anodal | 0.10 ± 0.16 | [-0.20; 0.41] |
| HD X Cathodal | -0.01 ± 0.14 | [-0.28; 0.27] |
| HD X Anodal | 0.14 ± 0.14 | [-0.15; 0.42] |
| **stimulationON X Cathodal** | **-0.32 ± 0.15** | **[-0.63; -0.03]** |
| stimulationON X Anodal | -0.03 ± 0.15 | [-0.33; 0.28] |
| TD X stimulationON X Cathodal | -0.18 ± 0.15 | [-0.49; 0.12] |
| TD X stimulationON X Anodal | -0.06 ± 0.15 | [-0.35; 0.23] |
| HD X stimulationON X Cathodal | 0.11 ± 0.13 | [-0.15; 0.39] |
| HD X stimulationON X Anodal | 0.00 ± 0.14 | [-0.26; 0.27] |
| Bayesian R<sup>2</sup> | 0.18 |  |

This table reports the results from a Bayesian hierarchical logistic regression model (equation (4)) explaining trial-wise health challenge success as a function of the categorical factor tDCS polarity (Anodal, Cathodal, and Sham), a dummy variable for session (stimulation=1, baseline=0), and continuous regressors for the standardized (z-score) absolute value of the healthiness difference (HD) and taste difference (TD) on each trial. The stimulationON regressor itself measures the difference between the initial baseline choice session and the second round of choices for the sham stimulation group (i.e. it captures potential effects of time and/or experience). Thus, the HD x stimulationON interaction indicates that the influence of healthiness on choice outcomes changes over time and/or experience with the food choice task. The effects of active tDCS are captured by the interaction coefficients, “stimulationON X Cathodal” and “stimulation X Anodal”. The model included subject-specific intercepts as well as subject-specific slopes for the effects of stimulation versus baseline (stimulationON), HD, and TD. Model fits are given as the population level mean of the posterior distribution ± standard deviation (SD) and the 95% credible interval.

**Supplementary Table 9.** Demographics of tDCS stimulation groups.

| Group | Sham | Cathodal | Anodal |
| --- | --- | --- | --- |
| BMI | 22.37 ± 2.75 | 22.30 ± 2.13 | 22.74 ± 2.92 |
| Age (in years) | 23.54 ± 2.90 | 23.75 ± 3.25 | 23.48 ± 23.04 |
| TFEQ – Restrained Eating | 9.94 ± 2.21 | 9.34 ± 2.46 | 9.59 ± 2.62 |
| TFEQ - Disinhibition | 7.52 ± 2.04 | 8 ± 2.17 | 7.70 ± 2.35 |
| TFEQ – Hunger susceptibility | 6.22 ± 2.11 | 6.43 ± 2.20 | 6.05 ± 2.26 |

This table reports descriptive statistics for Age, Body Mass Index (BMI) and scores on each subscale of the Three Factor Eating Questionnaire (TFEQ, German validated version by Pudel and Westenhöfer <sup>48</sup>). All values are given as mean and SD. Linear regression models testing for differences in each of these measures across stimulation groups did not reveal significant differences for any measure.

**Supplementary Table 10.** Mean and standard deviation for working memory by tDCS stimulation group (pre and post stimulation).

| Group | Mean pre | SD pre | Mean post | SD post |
| --- | --- | --- | --- | --- |
| Anodal | 7.39 | 1.45 | 7.84 | 1.53 |
| Cathodal | 7.18 | 1.23 | 7.61 | 1.29 |
| Sham | 7.12 | 1.29 | 7.38 | 1.31 |

The values in this table represent the forward digit span that we measured before and after stimulation using a computerized task that asked participants to remember sequences of 5 to 12 digits. Sequences were pseudo-randomized so that unique sequences would be presented before and after stimulation.

**Supplementary Table 11.** Comparison of working memory at baseline and after tDCS stimulation across stimulation groups.

| | Mean beta ± SD | 95% Credible Interval |
| --- | --- | --- |
| **Intercept** | **7.12 ± 0.18** | **[6.77; 7.47]** |
| Main Effect of stimulationON | 0.26 ± 0.25 | [-0.23; 0.73] |
| Cathodal Baseline | 0.05 ± 0.25 | [-0.45; 0.55] |
| Anodal Baseline | 0.27 ± 0.25 | [-0.23; 0.77] |
| Cathodal X stimulationON | 0.18 ± 0.36 | [-0.52; 0.86] |
| Anodal X stimulationON | 0.19 ± 0.36 | [-0.50; 0.91] |

**Supplementary Table 12.**
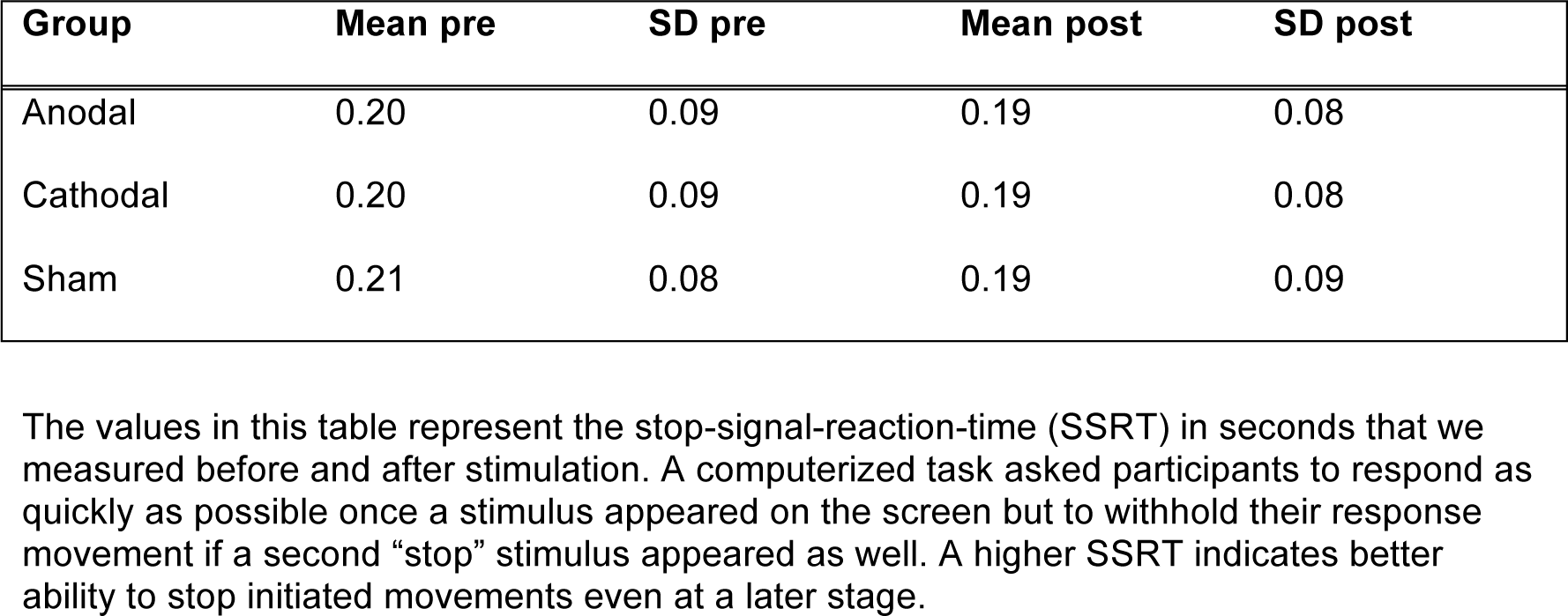
Mean and standard deviation for stop-signal-reaction-time (SSRT) in seconds by stimulation group (pre and post stimulation).

| Group | Mean pre | SD pre | Mean post | SD post |
| --- | --- | --- | --- | --- |
| Anodal | 0.20 | 0.09 | 0.19 | 0.08 |
| Cathodal | 0.20 | 0.09 | 0.19 | 0.08 |
| Sham | 0.21 | 0.08 | 0.19 | 0.09 |

The values in this table represent the stop-signal-reaction-time (SSRT) in seconds that we measured before and after stimulation. A computerized task asked participants to respond as quickly as possible once a stimulus appeared on the screen but to withhold their response movement if a second “stop” stimulus appeared as well. A higher SSRT indicates better ability to stop initiated movements even at a later stage.

**Supplementary Table 13.** Between-stimulation group comparison of stop signal reaction time (SSRT) at baseline and after stimulation.

| | Mean beta ± SD | 95% Credible Interval |
| --- | --- | --- |
| **Intercept** | **0.21 ± 0.01** | **[0.18; 0.24]** |
| Main Effect of stimulationON | -0.02 ± 0.02 | [-0.06; 0.02] |
| Cathodal Baseline | -0.01 ± 0.02 | [-0.05; 0.03] |
| Anodal Baseline | -0.01 ± 0.02 | [-0.05; 0.03] |
| Cathodal X stimulationON | 0.01 ± 0.03 | [-0.04; 0.06] |
| Anodal X stimulationON | 0 ± 0.03 | [-0.05; 0.06] |

**Supplementary Table 14.** Mean and standard deviation of intertemporal preference measures by stimulation group (pre and post stimulation).

| Group | Mean pre | SD pre | Mean post | SD post |
| --- | --- | --- | --- | --- |
| Anodal | 5123 | 1361 | 5214 | 1372 |
| Cathodal | 4628 | 1285 | 4583 | 1414 |
| Sham | 5144 | 1488 | 5123 | 1523 |

**Supplementary Table 15.**
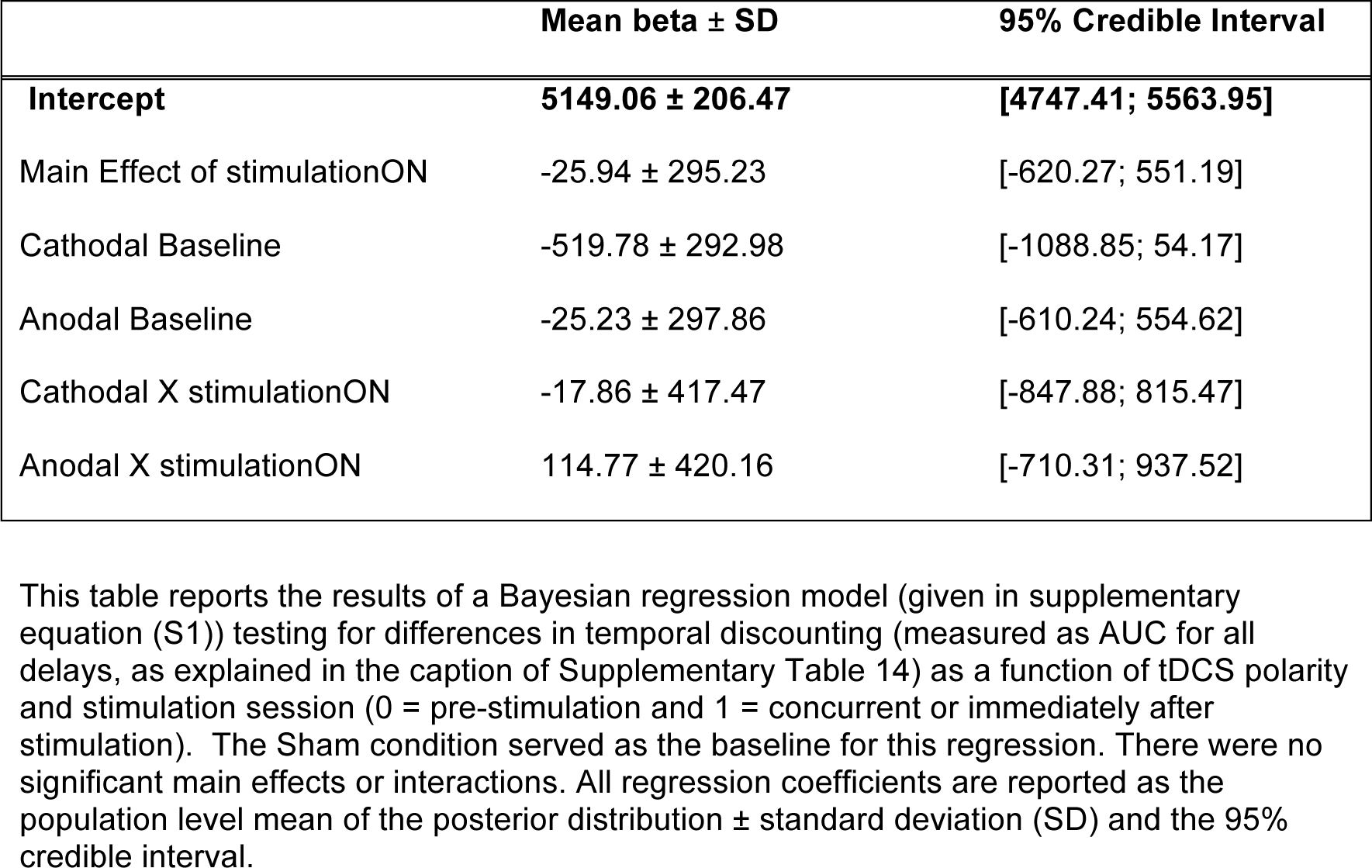
Between-stimulation group comparison of monetary intertemporal choice at baseline and after stimulation.

**Supplementary Table 16.**
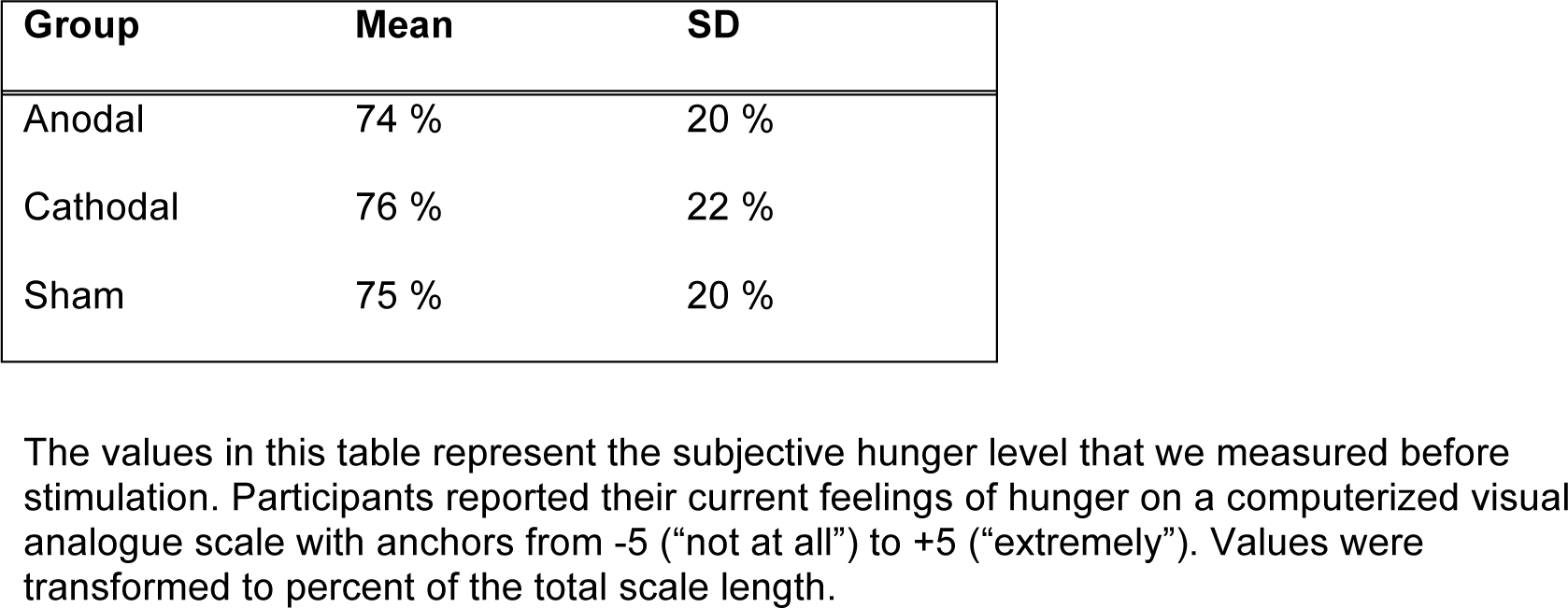
Hunger levels by tDCS stimulation group.

| Group | Mean | SD |
| --- | --- | --- |
| Anodal | 74 % | 20 % |
| Cathodal | 76 % | 22 % |
| Sham | 75 % | 20 % |

The values in this table represent the subjective hunger level that we measured before stimulation. Participants reported their current feelings of hunger on a computerized visual analogue scale with anchors from -5 (“not at all”) to +5 (“extremely”). Values were transformed to percent of the total scale length.

**Supplementary Table 17.** Baseline hunger level comparison between tDCS stimulation groups.

|  | Mean beta ± SD | 95% Credible Interval |
| --- | --- | --- |
| Intercept | 74.6 ± 2.73 | [69.17; 80.01] |
| Cathodal | 1.05 ± 3.90 | [-6.86; 8.57] |
| Anodal | -0.56 ± 3.93 | [-8.29; 7.1] |

**Supplementary Table 18.** Upper and lower bounds for each tDDM parameter during model fitting.

| Parameter | Lower bound | Upper bound |
| --- | --- | --- |
| $\omega_{\text{taste}}$ | -2 | 2 |
| $\omega_{\text{health}}$ | -2 | 2 |
| Thr | 0.6 | 3 |
| nDT | 0.01 | 1 |
| RST | -1 | 1 |
| bias | -1 | 1 |

Abbreviations:
$\omega_{\text{taste}}$ : weighting factor determining how much taste contributes to the evidence accumulation rate.
$\omega_{\text{health}}$ : weighting factor determining how much healthiness contributes to the evidence accumulation rate.
**Thr**: evidence threshold for responding (symmetric around zero)
**nDT**: non-decision time
**RST**: relative-start-time for health (positive values mean that health enters the process after taste, negative values mean health enters before taste)
**Bias**: starting point bias for the evidence accumulation process

**Supplementary Table 19.** The improvement in the log-likelihood for the tDDM relative to the standard DDM is proportional to increases in weighting parameters and relative-starting-times.

| Model parameter | Mean beta ± SD | 95% Credible Interval |
| --- | --- | --- |
| Intercept | 1.02 ± 0.59 | [-0.12; 2.14] |
| **\|$\omega_{\text{taste}}$\|** | **-0.45 ± 0.15** | **[-0.74; -0.17]** |
| **\|$\omega_{\text{health}}$\|** | **-0.62 ± 0.13** | **[-0.87; -0.36]** |
| **\|RST\|** | **-1.09 ± 0.20** | **[-1.49; -0.69]** |
| Thr | -0.27 ± 0.30 | [-0.85; 0.31] |
| nDT | 0.02 ± 0.48 | [-0.94; 0.97] |
| bias | -0.47 ± 0.28 | [-1.02; 0.06] |

This table lists the results from a Bayesian linear model (given in supplementary equation (S5)) explaining the difference in log-likelihood between the tDDM and the standard DDM by all parameters of the tDDM. In addition to threshold (Thr), non-decision time (nDT) and bias, absolute values of taste and health weights as well as relative-start-time (RST) were used in order to accommodate changes of these parameters in either direction. The standard DDM was fit using the same procedures as the tDDM, except that the equation omitted the relative-start-time parameter, which meant that both tastiness and healthiness entered into consideration at the same time. Note that similar results are obtained when using a logistic regression that treats the difference in log-likelihood as a binary outcome (tDDM better = 1, standard DDM better = 0).

**Supplementary Table 20.** A Bayesian linear regression testing for potential differences in the strength of the association between the relative attribute weights and starting times in generating versus recovered parameters.

| | Mean beta ± SD | 95% Credible Interval |
| --- | --- | --- |
| Intercept | 0.02 ± 0.03 | [-0.04; 0.07] |
| Relative weight | **0.15 ± 0.02** | **[0.11; 0.19]** |
| Recovered parameters | -0.03 ± 0.04 | [-0.10; 0.04] |
| Rel. weight* Rec. parameters | -0.02 ± 0.03 | [-0.07; 0.04] |
| Bayesian R<sup>2</sup> | 0.17 |  |

**Supplementary Table 21.**
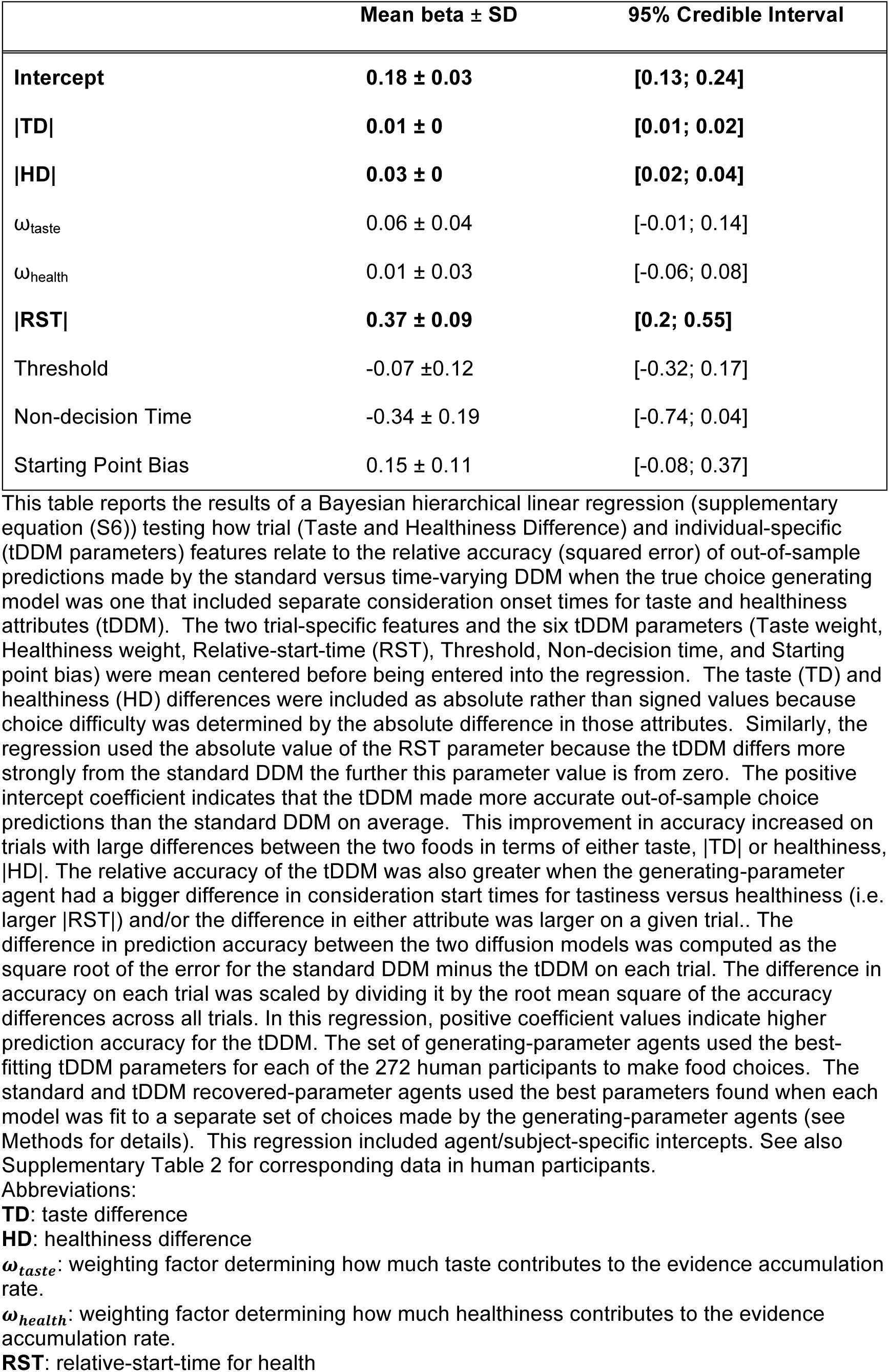
Improvement in out-of-sample prediction accuracy for the time-varying relative to the standard DDM on choices made by simulated agents using a time-varying DDM to make decisions.

| | Mean beta ± SD | 95% Credible Interval |
| --- | --- | --- |
| **Intercept** | **0.18 ± 0.03** | **[0.13; 0.24]** |
| **\|TD\|** | **0.01 ± 0** | **[0.01; 0.02]** |
| **\|HD\|** | **0.03 ± 0** | **[0.02; 0.04]** |
| $\omega_{\text{taste}}$ | 0.06 ± 0.04 | [-0.01; 0.14] |
| $\omega_{\text{health}}$ | 0.01 ± 0.03 | [-0.06; 0.08] |
| **\|RST\|** | **0.37 ± 0.09** | **[0.2; 0.55]** |
| Threshold | -0.07 ± 0.12 | [-0.32; 0.17] |
| Non-decision Time | -0.34 ± 0.19 | [-0.74; 0.04] |
| Starting Point Bias | 0.15 ± 0.11 | [-0.08; 0.37] |

**Supplementary Table 22.** Tests of differences in taste or health ratings across stimulation groups at baseline or following tDCS

| | Mean beta ± SD | 95% Credible Interval |
| --- | --- | --- |
| *A. Dependent variable = Health ratings* |  |  |
| **Intercept** (i.e. sham off) | **49.73 ± 2.03** | **[45.61; 53.68]** |
| **stimulationON** (i.e. sham on) | **1.03 ± 0.49** | **[0.07; 2.00]** |
| Cathodal | 0.74 ± 1.29 | [-1.77; 3.16] |
| Anodal | -1.18 ± 1.33 | [-3.65; 1.36] |
| Cathodal X stimulationON | -0.21 ± 0.69 | [-1.58; 1.13] |
| Anodal X stimulationON | -0.29 ± 0.69 | [-1.67; 1.11] |
| *B. Dependent variable = Taste ratings* |  |  |
| **Intercept** (i.e. sham off) | **52.85 ± 1.51** | **[49.93; 55.80]** |
| **stimulationON** (i.e. sham on) | **3.70 ± 0.95** | **[1.86; 5.57]** |
| **Cathodal** | **4.18 ± 1.92** | **[0.42; 7.91]** |
| **Anodal** | **4.92 ± 1.92** | **[1.14; 8.71]** |
| Cathodal X stimulationON | -0.16 ± 1.31 | [-2.75; 2.41] |
| Anodal X stimulationON | -1.07 ± 1.31 | [-3.69; 1.49] |

This table reports the results from two hierarchical Bayesian regressions (given in supplementary equation (S3)) testing for differences in taste or health ratings across stimulation groups at baseline or following tDCS. Taste and health ratings were coded as percent maximum tastiness or healthiness. Both regressions included regressors for the main effect of stimulation group (sham, anodal, cathodal), stimulation session (baseline, post-stimulation), and their interaction as population-level effects. The models also included intercepts for subjects and food items (i.e., each subject and food item was treated as a random effect) as well as subject-specific and food-specific slopes for the effect of stimulation versus baseline. **A)** Health ratings increased at the second rating during or after stimulation, but no group differences were observed at baseline or following stimulation. **B)** Taste ratings were higher at baseline for the anodal and cathodal groups compared to the sham group and increased for all groups at the second rating. Critically, there was no stimulation group X stimulation session interaction, indicating that stimulation condition did not differentially influence the ratings at the second stimulation session.

**Supplementary Table 23.** Effects of the average taste and health rating response times on the relative-start-time for health.

| | Mean beta ± SD | 95% Credible Interval |
| --- | --- | --- |
| **Intercept** | **-0.63 ± 0.25** | **[-1.12; -0.13]** |
| Mean RT health | 0.05 ± 0.11 | [-0.17; 0.28] |
| Mean RT taste | 0.07 ± 0.12 | [-0.17; 0.31] |
| nDT | 0.33 ± 0.24 | [-0.13; 0.80] |

This table reports the results from a Bayesian regression (see supplementary equation (S4)) of the tDDM-estimated relative-start-time for health in baseline choices on the rating speed for taste and health, in the sample of participants from the TDCS study. Model fits are given as the population level mean of the posterior distribution ± standard deviation (SD) and the 95% credible interval. There was no significant relation between the mean reaction time (RT) for health and taste ratings (in seconds) and the relative-start-time for health in the individual choice process.

**Supplementary Table 24.** Bayesian logistic regression for healthy choice outcomes as a function of attribute weighting and timing parameters.

| | Mean beta ± SD | 95% Credible Interval |
| --- | --- | --- |
| Intercept | -0.17 ± 0.26 | [-0.68; 0.35] |
| $\omega_{\text{taste}}$ | **0.52 ± 0.17** | **[0.18; 0.85]** |
| $\omega_{\text{health}}$ | **1.43 ± 0.21** | **[1.02; 1.87]** |
| **RST** | **0.72 ± 0.36** | **[0.01; 1.43]** |
| **Challenge** | **-0.73 ± 0.25** | **[-1.22; -0.25]** |
| **Taste Difference (TD)** | **0.19 ± 0.03** | **[0.14; 0.24]** |
| **Health Difference (HD)** | **0.69 ± 0.08** | **[0.54; 0.84]** |
| $\omega_{\text{taste}}$ X RST | -0.29 ± 0.27 | [-0.81; 0.23] |
| $\omega_{\text{health}}$ X RST | **-1.34 ± 0.26** | **[-1.86; -0.85]** |
| $\omega_{\text{taste}}$ X Challenge | **-1.60 ± 0.18** | **[-1.95; -1.26]** |
| $\omega_{\text{health}}$ X Challenge | -0.07 ± 0.21 | [-0.50; 0.34] |
| RST X Challenge | -0.23 ± 0.38 | [-0.96; 0.49] |
| $\omega_{\text{taste}}$ X RST X Challenge | -0.28 ± 0.27 | [-0.83; 0.24] |
| $\omega_{\text{health}}$ X RST X Challenge | 0.22 ± 0.27 | [-0.26; 0.78] |

This table reports results from a Bayesian logistic regression (given in supplementary equation (S8)) modelling healthy food choices as a function of the estimated taste and health weighting parameters of the tDDM as well as the relative start time for health (RST). The model also included the signed difference in taste (TD) and healthiness (HD) ratings between the healthier and less healthy option (i.e. healthier – less healthy), and a dummy variable (Challenge) indicating whether participants faced a challenge on the respective trial. The model included subject-specific intercepts as well as subject-specific slopes for the TD, HD and Challenge regressors. Data were pooled across the baseline conditions of all 4 studies. Model fits are given as the population level mean of the posterior distribution ± standard deviation (SD) and the 95% credible interval. Abbreviations:
$\omega_{\text{taste}}$ : weighting factor determining how much taste contributes to the evidence accumulation rate.
$\omega_{\text{health}}$ : weighting factor determining how much healthiness contributes to the evidence accumulation rate.
**RST**: relative-start-time for health

**Supplementary Table 25.** Bayesian linear regression for response times in health challenge and non-challenge trials as a function of tDDM parameters.

| | Mean beta ± SD | 95% Credible Interval |
| --- | --- | --- |
| **Intercept** | **0.55 ± 0.04** | **[0.47; 0.63]** |
| **Challenge** | **-0.12 ± 0.04** | **[-0.19; -0.05]** |
| $\omega_{\text{taste}}$ | -0.04 ± 0.03 | [-0.11; 0.02] |
| $\omega_{\text{health}}$ | **-0.09 ± 0.04** | **[-0.16; -0.02]** |
| RST | -0.07 ± 0.07 | [-0.21; 0.06] |
| **Healthy choice (HC)** | **-0.07 ± 0.03** | **[-0.14; -0.01]** |
| **\| Taste Difference (TD) \|** | **-0.06 ± 0.01** | **[-0.07; -0.04]** |
| **\| Health Difference (HD) \|** | **-0.02 ± 0.01** | **[-0.03; -0.01]** |
| $\omega_{\text{taste}}$ X RST | 0.08 ± 0.05 | [-0.03; 0.18] |
| $\omega_{\text{health}}$ X RST | 0.07 ± 0.05 | [-0.03; 0.15] |
| $\omega_{\text{taste}}$ X HC | 0.02 ± 0.03 | [-0.03; 0.08] |
| $\omega_{\text{health}}$ X HC | -0.01 ± 0.03 | [-0.07; 0.05] |
| RST X HC | 0.03 ± 0.05 | [-0.07; 0.13] |
| Challenge X $\omega_{\text{taste}}$ | 0.03 ± 0.03 | [-0.03; 0.09] |
| **Challenge X $\omega_{\text{health}}$** | **0.08 ± 0.03** | **[0.01; 0.14]** |
| Challenge X RST | 0.06 ± 0.06 | [-0.06; 0.18] |
| Challenge X HC | 0.07 ± 0.04 | [0; 0.13] |
| **Challenge X \|TD\|** | **0.05 ± 0.01** | **[0.03; 0.06]** |
| Challenge X \|HD\| | 0 ± 0.01 | [-0.01; 0.02] |
| $\omega_{\text{taste}}$ X RST X HC | -0.07 ± 0.04 | [-0.16; 0.01] |
| $\omega_{\text{health}}$ X RST X HC | 0.05 ± 0.04 | [-0.03; 0.12] |
| **Challenge X $\omega_{\text{taste}}$ X RST** | **-0.11 ± 0.05** | **[-0.21; -0.02]** |
| Challenge X $\omega_{\text{health}}$ X RST | -0.06 ± 0.04 | [-0.14; 0.02] |
| Challenge X $\omega_{\text{taste}}$ X HC | 0 ± 0.03 | [-0.06; 0.07] |
| Challenge X $\omega_{\text{health}}$ X HC | -0.04 ± 0.03 | [-0.11; 0.02] |
| Challenge X RST X HC | -0.04 ± 0.06 | [-0.15; 0.08] |
| **Challenge X $\omega_{\text{taste}}$ X RST X HC** | **0.17 ± 0.05** | **[0.07; 0.26]** |
| Challenge X $\omega_{\text{health}}$ X RST X HC | 0.04 ± 0.04 | [-0.05; 0.12] |

This table reports results from a Bayesian hierarchical linear regression (given in supplementary equation (S9)) modelling response times in food choices as a function of the estimated taste and health weighting parameters of the tDDM as well as the relative start time for health (RST), and the absolute distance in taste (|TD|) and health (|HD|) ratings between the two options. Dummy variables indicated whether participants faced a challenge on the respective trial and whether they chose the healthier food (HC). The model included subject-specific intercepts and subject-specific slopes for the |TD| and |HD| regressors as well as the interactions of these regressors with the challenge trial and healthy choice dummy variables to capture individual differences in sensitivity to taste and health aspects when making healthy or unhealthy choices in challenging or non-challenging settings. Data were pooled across the baseline conditions of all 4 studies. Model fits are given as the population level mean of the posterior distribution ± standard deviation (SD) and the 95% credible interval. Abbreviations:
$\omega_{\text{taste}}$ : weighting factor determining how much taste contributes to the evidence accumulation rate.
$\omega_{\text{health}}$ : weighting factor determining how much healthiness contributes to the evidence accumulation rate.
RST: relative-start-time for health

**Supplementary Table 26.** Bayesian linear regression for response times in cases where the healthier option was chosen.

| | Mean beta ± SD | 95% Credible Interval |
| --- | --- | --- |
| **Intercept** | **0.41 ± 0.02** | **[0.37; 0.46]** |
| **\| Taste Difference (TD) \|** | **-0.09 ± 0.01** | **[-0.11; -0.06]** |
| **\| Health Difference (TD) \|** | **-0.02 ± 0.01** | **[-0.04; 0.00]** |
| Challenge | -0.03 ± 0.02 | [-0.07; 0.02] |
| **\|TD\| X Challenge** | **0.12 ± 0.01** | **[0.10; 0.15]** |
| **\|HD\| X Challenge** | **-0.04 ± 0.01** | **[-0.07; -0.02]** |

This table reports results from a Bayesian linear regression (given in supplementary equation (S10)) modelling response times in trials where participants selected the healthier option as a function the absolute difference in taste (|TD|) and health (|HD|) ratings between the two options. A dummy variable (challenge) indicated whether participants faced a health challenge on the respective trial because the healthier option was not the more tasty option. The model included subject-specific intercepts and subject-specific slopes for the |TD| and |HD| regressors and their interaction with the challenge trial dummy variable to capture individual differences in sensitivity to taste and health aspects in challenging or non-challenging settings. Data were pooled across the baseline conditions of all 4 studies. Model fits are given as the population level mean of the posterior distribution ± standard deviation (SD) and the 95% credible interval. Participants' response times were associated with the differences in taste between the two options when they chose the healthier option in both health challenge and non-challenge trials.

**Supplementary Table 27.** Bayesian linear regression for response times in cases where the tastier option was chosen.

| | Mean beta ± SD | 95% Credible Interval |
| --- | --- | --- |
| **Intercept** | **0.45 ± 0.02** | **[0.41; 0.49]** |
| **\| Taste Difference (TD) \|** | **-0.08 ± 0.01** | **[-0.10; -0.06]** |
| **\| Health Difference (TD) \|** | **-0.02 ± 0.01** | **[-0.03; -0.00]** |
| **Challenge** | **-0.06 ± 0.02** | **[-0.09; -0.02]** |
| **\|TD\| X Challenge** | **0.04 ± 0.01** | **[0.02; 0.06]** |
| **\|HD\| X Challenge** | **0.05 ± 0.01** | **[0.03; 0.07]** |

This table reports results from a Bayesian linear regression (given in supplementary equation (S10)) modelling response times in trials where participants selected the tastier option as a function the absolute difference in taste (|TD|) and health (|HD|) ratings between the two options. A dummy variable (challenge) indicated whether participants faced a health challenge on the respective trial because the healthier option was not the more tasty option. The model included subject-specific intercepts and subject-specific slopes for the |TD| and |HD| regressors and their interaction with the challenge trial dummy variable to capture individual differences in sensitivity to taste and health aspects in challenging or non-challenging settings. Data were pooled across the baseline conditions of all 4 studies. Model fits are given as the population level mean of the posterior distribution ± standard deviation (SD) and the 95% credible interval. Participants' response times were associated with the differences in healthiness between the two options when they chose the tastier option in both health challenge and non-challenge trials.

**Supplementary Table 28.**
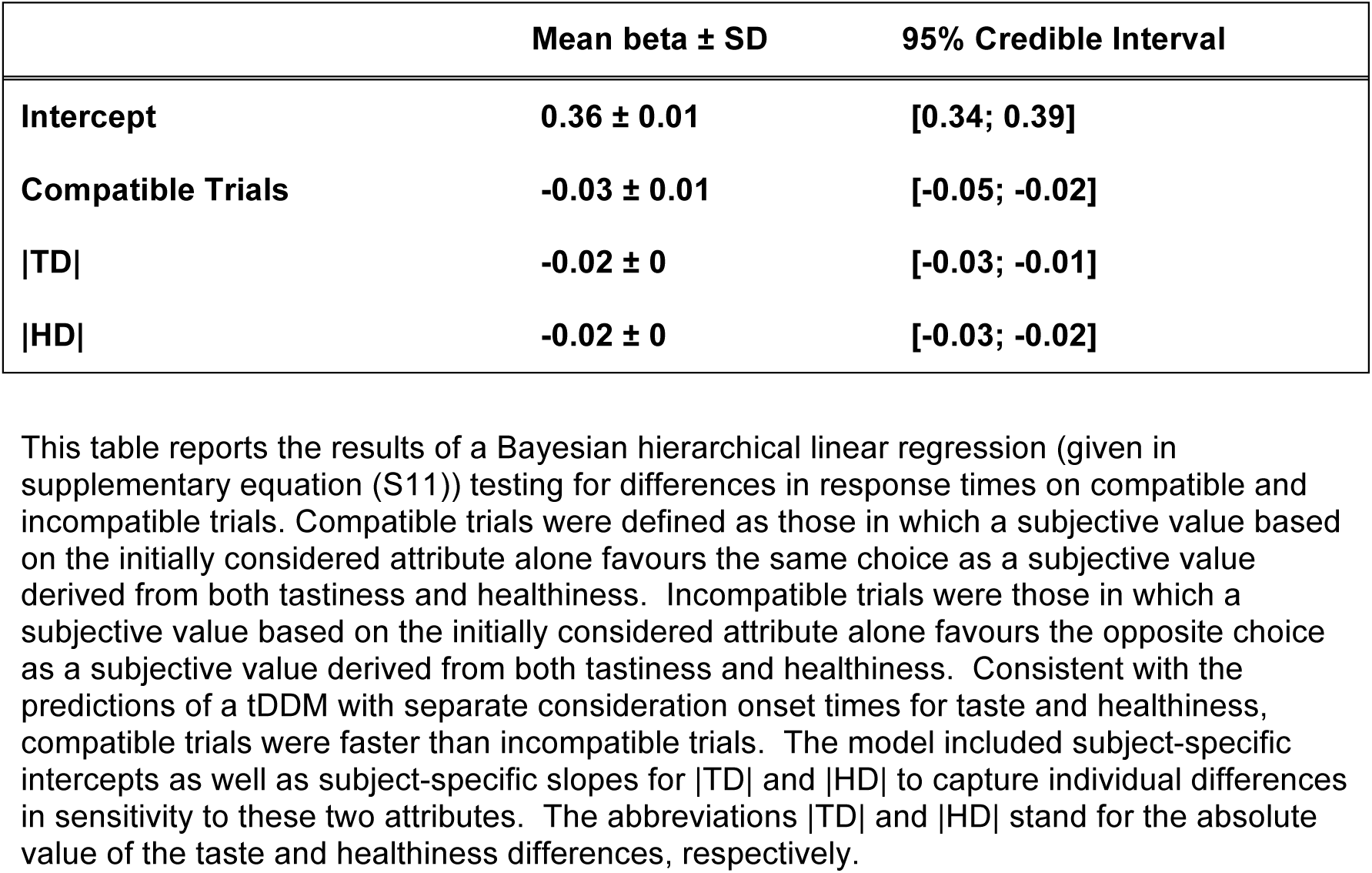
Response times as a function of whether a subjective value based on the initially considered attribute alone favours the same choice as a subjective value derived from both tastiness and healthiness.

| | Mean beta ± SD | 95% Credible Interval |
| --- | --- | --- |
| **Intercept** | **0.36 ± 0.01** | **[0.34; 0.39]** |
| **Compatible Trials** | **-0.03 ± 0.01** | **[-0.05; -0.02]** |
| **\|TD\|** | **-0.02 ± 0** | **[-0.03; -0.01]** |
| **\|HD\|** | **-0.02 ± 0** | **[-0.03; -0.02]** |

This table reports the results of a Bayesian hierarchical linear regression (given in supplementary equation (S11)) testing for differences in response times on compatible and incompatible trials. Compatible trials were defined as those in which a subjective value based on the initially considered attribute alone favours the same choice as a subjective value derived from both tastiness and healthiness. Incompatible trials were those in which a subjective value based on the initially considered attribute alone favours the opposite choice as a subjective value derived from both tastiness and healthiness. Consistent with the predictions of a tDDM with separate consideration onset times for taste and healthiness, compatible trials were faster than incompatible trials. The model included subject-specific intercepts as well as subject-specific slopes for |TD| and |HD| to capture individual differences in sensitivity to these two attributes. The abbreviations |TD| and |HD| stand for the absolute value of the taste and healthiness differences, respectively.

